# The evolutionary ancient MEIS transcription factors actuate lineage-specific transcription to establish cardiac fate

**DOI:** 10.1101/2024.07.18.604075

**Authors:** Zoulfia Darieva, Peyman Zarrineh, Naomi Phillips, Joshua Mallen, Araceli Garcia Mora, Ian Donaldson, Laure Bridoux, Megan Douglas, Sara F Dias Henriques, Dorothea Schulte, Matthew J Birket, Nicoletta Bobola

## Abstract

Control of gene expression is commonly mediated by distinct combinations of transcription factors (TFs). This cooperative action allows multiple biological signals to be integrated at specific regulatory elements, resulting in highly specific gene expression patterns in space and time. It is unclear whether combinatorial binding is also necessary to bring together TFs with distinct biochemical functions, which collaborate to effectively recruit and activate RNA polymerase II. Using a cardiac differentiation model, we find that the largely ubiquitous, evolutionary ancient homeodomain proteins MEIS are essential for activating a cardiac-specific gene expression program. MEIS TFs act as *actuators*, fully activating transcriptional programs selected by lineage-restricted TFs to drive the dynamic progression of cardiac differentiation. Combinatorial binding of MEIS with lineage-enriched TFs, GATA and HOX, provides selectivity, guiding MEIS to function at cardiac-specific enhancers. In turn, MEIS TFs promote accumulation of the methyltransferase KMT2D to initiate lineage-specific enhancer commissioning. MEIS combinatorial binding dynamics, dictated by the changing dosage of its partners, drive cells into progressive stages of cardiac differentiation. Our results uncover tissue-specific transcriptional activation as the result of ubiquitous *actuator TFs* harnessing general transcriptional coactivators at tissue-specific enhancers, to which they are directed by binding with lineage- and domain-specific TFs.

## Introduction

Control of gene expression determines and maintains cellular identity and function in embryonic development and adult tissue homeostasis. Lineage-specific TFs orchestrate the precise gene expression programs that determine cell fate [1]. However, with rare exceptions, lineage-specific TFs are unable to efficiently program cells into their specific cell type [2]. Examples from animal development or cellular reprogramming indicate that transcriptional regulation is commonly mediated by distinct combinations of TFs [3]. Combinatorial binding enables the integration of multiple biological inputs at cis-regulatory elements to generate highly specific regulatory outcomes in space and time [1]. This collaborative binding is favoured because multiple TFs, interacting with DNA simultaneously, display higher affinity for their designated sites [4,5]. The role of TF cooperativity in stabilizing TF-DNA binding is well-established. However, our understanding of how these TF combinations regulate RNA polymerase II activity to affect gene expression remains incomplete. The biochemical effects of TFs after binding DNA are largely unmapped. Specifically, it is unclear if DNA-bound TFs recruit co-activators equivalently, making them interchangeable, or if combinatorial binding assembles TF complexes with emerging functional properties crucial for transcriptional activation.

MEIS TFs belong to the three amino acid loop extension (TALE) superclass, which includes evolutionary ancient TFs with an atypical homeodomain [6-8]. Initially discovered as HOX partners, MEIS1-2 TFs are essential for development of many organs and can function as oncogenes [9]. While largely ubiquitous, MEIS1-2 TFs have been implicated in the control of tissue-specific transcription. In mouse development, high-confidence MEIS binding events occur at tissue-specific locations across different embryonic tissues. These binding events are enriched in motifs recognised by tissue-specific TFs and associated with increased enhancer activity and gene expression in the same tissue [10,11]. The potential of widely expressed TFs like MEIS to drive the acquisition of tissue-specificity, and the precise mechanisms by which they might achieve this, remain an open question, with implications for cell reprogramming.

Here we find that MEIS proteins act as *actuator TFs*, fully activating the transcriptional programs directed by lineage-specific TFs to orchestrate the dynamic progression of cardiac differentiation. MEIS recognition motifs are enriched in human developmental enhancers active in distinct tissues, heart, brain, limb, kidney and adrenal glands. Using a cardiac differentiation system, we find that MEIS is essential to initiate a cardiac-specific gene expression program. Combinatorial binding of MEIS with GATA and HOX directs MEIS to function at cardiac-specific enhancers. Here, MEIS promotes accumulation of the methyltransferase KMT2D to initiate lineage-specific enhancer commissioning. MEIS trades partners to orchestrate the dynamic progression of cardiac differentiation. Our results uncover tissue-specific transcriptional activation as the result of ubiquitous actuator TFs harnessing general transcriptional activator at tissue-specific enhancers, to which they are directed by binding with lineage- and domain-specific TFs.

## Results

### MEIS1-2 are essential for cardiac lineage differentiation

Using a screen designed to map TF combinatorial binding in human developmental enhancers [12], we found a significant enrichment of recognition motifs for TALE TFs nearby tissue-specific sequence signatures. TALE motifs occur in proximity of sequences recognized by tissue-restricted TFs in heart (ventricle), brain, upper limb, adrenal gland and kidney enhancers (Fig 1A). In contrast to their potential co-binding partners, MEIS TFs are broadly expressed (Fig 1B). Cardiac TFs are the most highly significant predicted partners of MEIS. Therefore, to understand the exact contribution of ubiquitous TFs like MEIS to to the acquisition of tissue-specificity, we exploited a well-established 3D model of cardiac differentiation (Fig. 1C) [13]. Initially, we investigated the expression dynamics of *MEIS* genes within this system. Single cell analysis of human embryonic stem cells (hESC) differentiation into cardiomyocytes (Fig 1D) reflects the temporal progression from mesodermal cells, marked by *TBXT* at d3, to *MESP1*-positive cardiac mesoderm (Fig 1E). The expression of *MEIS1* and *MEIS2* follows cardiac mesoderm induction and coincides with the emergence of cardiac progenitor cells (Fig 1F). Both *MEIS1* and *MEIS2* remain highly expressed in *MEF2C*-*TBX5*-positive late progenitors (Fig. 1G). While cardiomyocytes (identified by the expression of *NKX2-5* and *MYH6*) (Fig 1H) and epicardial cells (*WT1*-*TBX18*-positive) (Fig 1I) maintain high *MEIS2* expression, *MEIS1* expression decreases at later, differentiated stages (Fig. 1F; Fig. 1J). Relative to *MEIS1* and *MEIS2*, whose expression is first detected in early progenitors, *MEIS3* is already expressed in the mesoderm and exhibits less pronounced temporal dynamics throughout differentiation (Fig S1A). Likewise, *PBX* genes, whose encoded TFs also bind the TALE motif, are expressed consistently across the analyzed stages of differentiation (Fig S1B).

**Figure 1.**
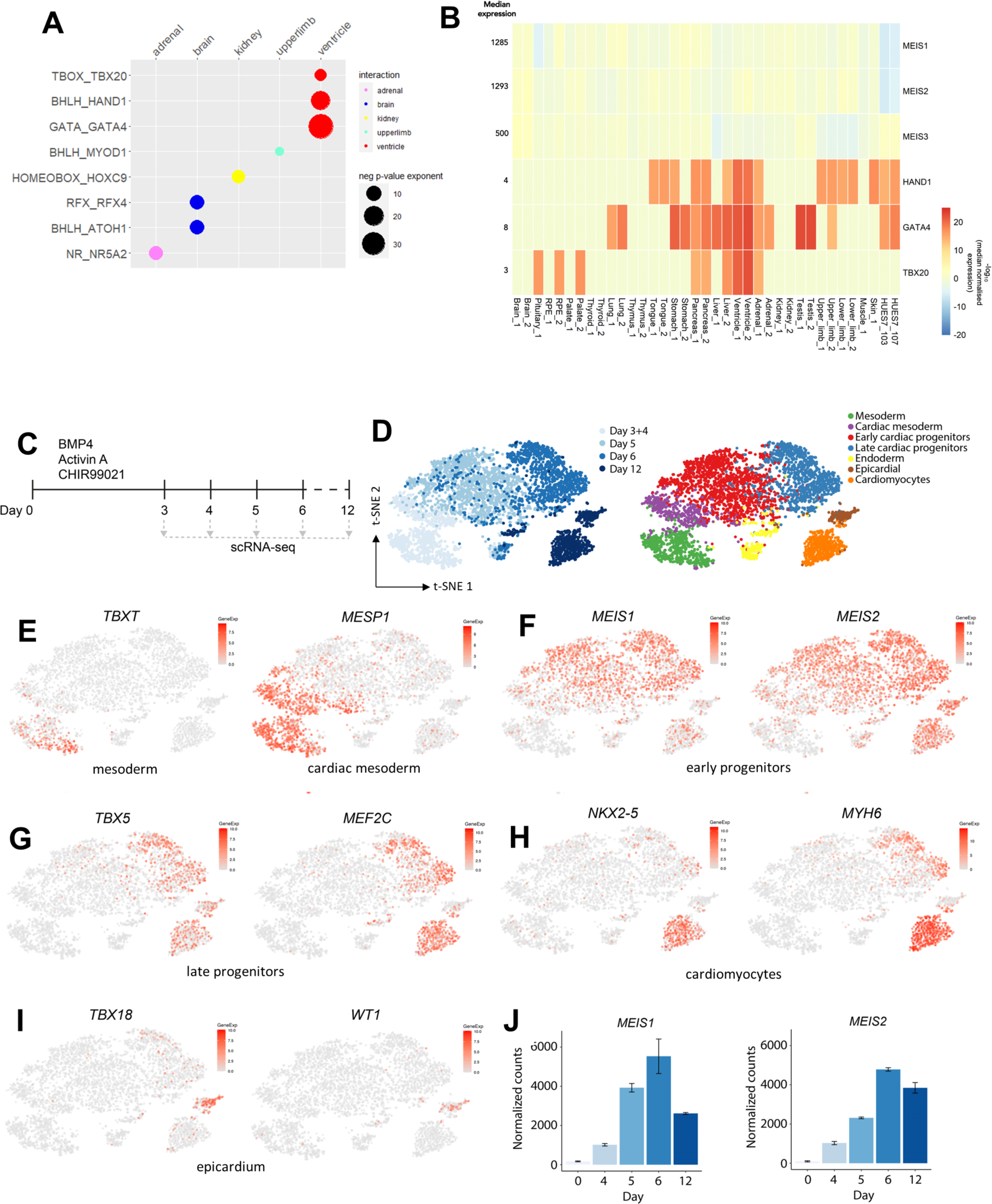
Tissue-specific patterns of ubiquitous TFs. A. TALE binding site co-occurs with motifs recognised by tissue-restricted TFs at enhancers active in human embryonic development data sourced from [56]. The tissue-restricted TFs listed on the left are predicted to bind signature motifs enriched in human enhancers specifically active in the tissue shown (top). TALE motif was identified as enriched within 100nt upstream or downstream of the tissue-restricted TF motif. A cut-off of p < 1x10^-5^ and enrichment in > 5% of foreground regions were used. The size of the bubbles matches the negative log p-value: larger bubbles represented smaller p-values. B. Expression of MEIS1-2 and their predicted cardiac partners in human embryonic tissues; data sourced from [56]. All tissues are duplicated except for pituitary, muscle and skin. Human embryonic RNA-seq read raw counts were down sampled based on the 75th percentile and read counts for each gene were divided by the gene’s median read counts. Negative logarithm of the median normalised counts were used to plot the heatmap. C. Schematic of cardiac differentiation protocol of hESCs and collection timepoints for scRNA-seq. D. t-SNE plots of scRNA-seq cardiac differentiation time-course coloured by day and cell type. E-I. t-SNE plots showing gene expression level of key markers for mesoderm (E), early (F) and late (G) progenitors, cardiomyocytes (H) and epicardial cells (I). Relative to the progression of differentiation, the earliest *MEIS1-* and *MEIS2-* positive cells correspond to early progenitors. J. Bulk RNA-seq normalized counts for MEIS1 and MEIS2 of hESCs undergoing cardiac differentiation.

*MEIS1-2* expression in early cardiac progenitors is suggestive of a role in cardiac lineage determination. Starting from a NKX2-5-GFP reporter hESC line [14], we used CRISPR Cas9 to generate a knock-out of MEIS1 and MEIS2 (referred to as MEIS KO) (Fig S2A). Wild-type embryoid bodies (EBs) became GFP-positive by day (d) 8 of differentiation and contracted soon after, indicating proper cardiomyocyte differentiation; in contrast, MEIS KO EBs remained GFP-negative and failed to contract (Fig 2A). At d12, MEIS KO EBs lacked markers of cardiomyocytes and epicardial cells, indicating a general failure in cardiac progenitor specification or differentiation (Fig S2B). The absence of MEIS1-2 in KO hESC did not affect the expression of pluripotency factors or markers of mesoderm differentiation (*TBXT*, *EOMES*, *MESP1*), indicating unaffected initial stages (Fig 2B). However, dysfunction was evident by the failed upregulation of crucial cardiac TFs like *GATA4* and *NKX2-5* in the absence of MEIS1-2 (Fig 2B).

**Figure 2.**
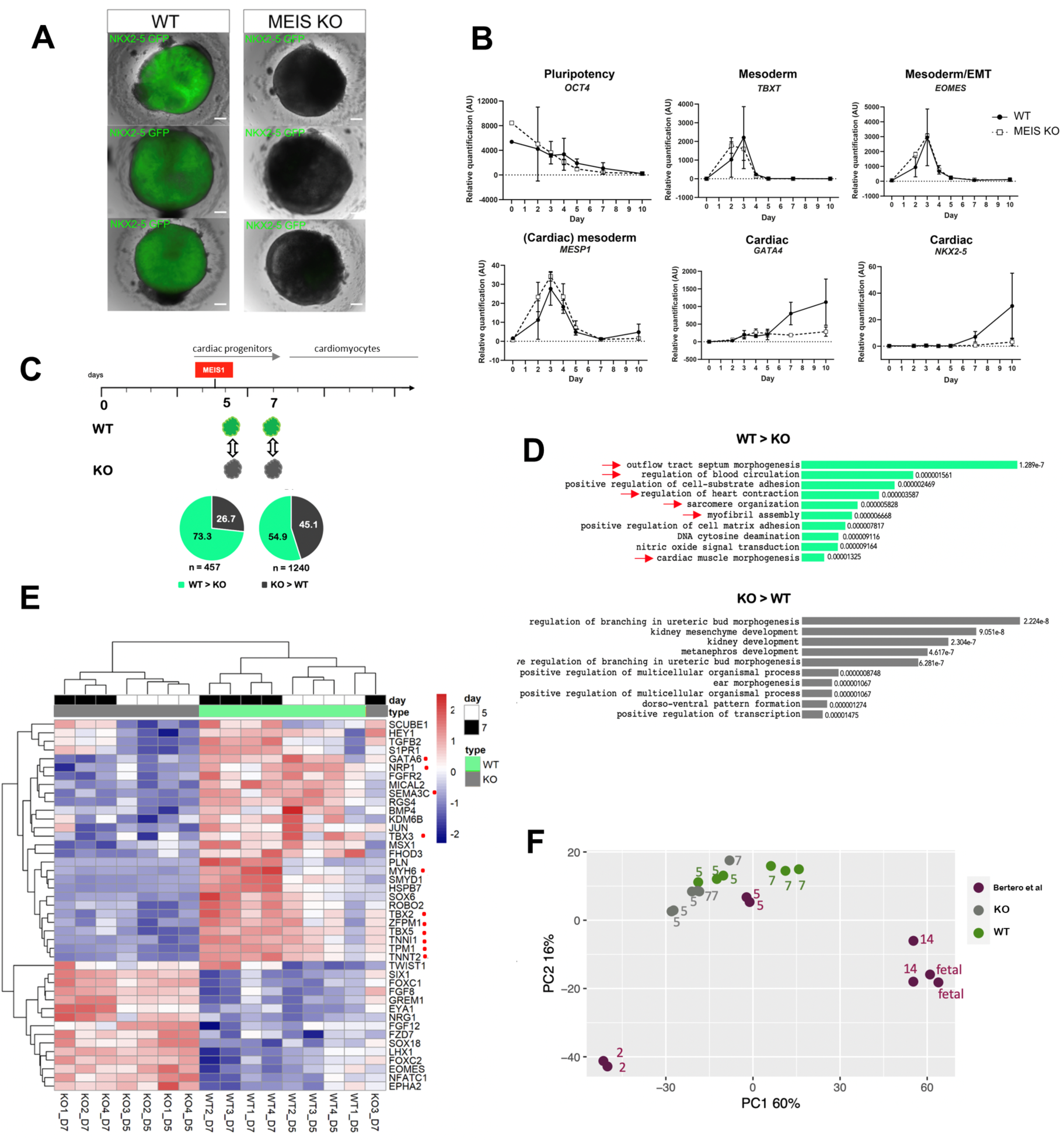
MEIS1-2 are essential for cardiac differentiation. A. Live imaging of cardiac lineage reporter *NKX2-5-GFP* in wild-type and MEIS KO EBs at d12. In the same conditions, MEIS KO EBs fail to activate *NKX2-5* expression. The GFP channel is overlayed on the brightfield images. B. Relative gene expression of pluripotency, mesoderm and cardiac mesoderm markers, measured by qPCR, in wild-type and MEISKO hESC undergoing cardiac differentiation. AU = arbitrary units. C. Schematic of the comparative analysis of wild type and MEIS KO transcriptomes with the proportion of up- and down-regulated genes at d5 and d7 of cardiac differentiation. The onset of *MEIS1* activation is indicated above the timeline. D. GO associated with genes down- and up-regulated in the absence of MEIS1-2. Genes downregulated in the MEIS KO (green) are almost exclusively associated with cardiovascular terms while upregulated genes (dark grey) display a variety of developmental GO terms, none associated with cardiac development. Only the top 10 GO terms are shown. E. Changes in expression of cardiac mesoderm GO genes in wild type (green) and MEIS KO (grey) cells at d5 (white) and d7(black). Red dots indicate genes with a cardiac phenotype or cardiac-specific expression. F. PCA plots comparing human cardiac differentiation (d2, d5, d14) and fetal heart (fetal) transcriptomes (purple), sourced from [15] alongside d5 and d7 wild-type (green) and MEIS KO (grey) transcriptomes. D5-7 MEIS KO cells cluster more closely than wild-type cells along PC1 (representing time), indicating a disruption in the trajectory toward cardiac differentiation.

To examine the requirement for MEIS1-2 in cardiomyocyte differentiation, we compared transcriptomes of wild-type and MEIS KO at two stages, corresponding to early (d5, shortly after the onset of *MEIS1* transcription) and late (d7) cardiac progenitors. We found a total of 457 and 1240 differentially expressed (DE) genes, respectively (log fold change > [1]; p <0.05), at d5 and d7, respectively) (Fig 2C; Table S1). At the earliest time point, downregulated genes in the KO represent the largest fraction of dysregulated genes (>70%). By d7, there is a similar ratio of upregulated and downregulated genes. Changes measured close to the start of MEIS expression (d5) likely reflect direct regulation by MEIS, suggesting that MEIS transcription factors predominantly function as transcriptional activators. Genes activated by MEIS (d5 down in KO) are associated with heart development and cardiac differentiation. In contrast, upregulated genes (d5 up in KO) are linked to multiple embryonic processes, but not including cardiac development (Fig 2D). When genes associated with cardiac mesoderm gene ontologies (GO) are selected, genes downregulated in MEIS KO are enriched in cardiac-specific transcripts, while upregulated genes are generally expressed in mesoderm (e.g. *TWIST1*, *FOXC1-2*, *SIX1*) (Fig 2E). These results indicate that MEIS KO cells fail to activate cardiac-specific genes, possibly remaining in a more primitive mesodermal stage. Consistent with this hypothesis, PCA analysis positions MEIS KO cells between mesoderm and cardiac progenitors in a time course of cardiac differentiation [15], with limited change in MEIS KO cells from d5 to d7 (Fig 2F). In sum, these findings indicate that MEIS1-2 are essential TFs for cardiac lineage differentiation, highlighting the requirement of ubiquitous TFs in tissue-specific transcription.

### MEIS1-2 function as activators to establish a cardiac-specific program

Physical access to DNA is a dynamic property of chromatin that reflects the cis-regulatory control of gene expression. To examine the effects of MEIS on the chromatin landscape, we mapped chromatin accessibility in wild-type and MEIS KO cells and assessed quantitative differences in ATAC-seq signal intensities at early (d5) and late (d12) cardiac differentiation stages. Globally, MEIS1-2 are (directly or indirectly) responsible for a relatively small fraction of chromatin accessibility: more than 70% open chromatin remains unperturbed in MEIS KO cells and only around 15% regions loose accessibility and an equal fraction gains accessibility (Fig 3A). Strikingly, however, the fraction of regions whose gain in accessibility depends on MEIS (ATAC WT>KO), is specifically associated with cardiovascular development and differentiation (Fig 3B). Chromatin accessibility shows distinct temporal dynamics: at d5, higher accessibility is largely contained in distal intergenic and intronic regions, while at d12, promoters account for half of the regions that increase in accessibility in the presence of MEIS. In contrast, regions whose accessibility increases in the absence of MEIS, are located in intergenic and intronic regions at both time points (Fig. S3A). This discrepancy likely reflects the failure of mutant cells to progress to a fully differentiated phenotype, characterized by enhancer decommissioning and consequent loss of accessibility at distal and intronic regions [16]. To identify chromatin and transcriptional changes initiated by MEIS, we mapped MEIS1 genomic occupancy in cardiac progenitors (d5), shortly after *MEIS1* expression. High-confidence MEIS1 peaks (FE>10; n=1742; Table S2) largely occur distally from any known transcriptional start site (TSS), with only a small fraction of MEIS binding (<5%) in the vicinity (within 5 kb) of a TSS (Fig S3B). GREAT [17] largely associated MEIS1 peaks to genes involved in heart development (Fig 3C). Chromatin accessibility and acetylation are hallmark features of active chromatin [18,19]. The top fraction of MEIS1 peaks almost exclusively intersect regions exhibiting increased accessibility in wild-type cells (Fig S3C). At the onset of differentiation (d5), we observed a significant reduction in chromatin accessibility (Fig 3D) and acetylation (Fig 3E) in MEIS KO cells; this shift is specific to regions normally occupied by MEIS1. The *PRDM6* and *TBX5* loci exemplify loss of chromatin accessibility and acetylation in the absence of MEIS (Fig 3F). Finally, we assigned MEIS1 peaks to genes and intersected genes associated with MEIS1 binding with DE genes at the same time point (d5). MEIS directly activates 133 genes (out of a total of 335 downregulated in the mutant) and repress 20 genes (out of a total of 122 upregulated in the mutant) (Fig 3G). In sum, these results indicate that MEIS TFs predominantly function as transcriptional activators, promoting chromatin accessibility and acetylation at critical regions for cardiac differentiation, and the expression of their associated genes. The enrichment of intrinsically disordered regions (IDRs) in MEIS1 and MEIS2 also supports their role in activating transcription [20,21] (Fig S3DE).

**Figure 3.**
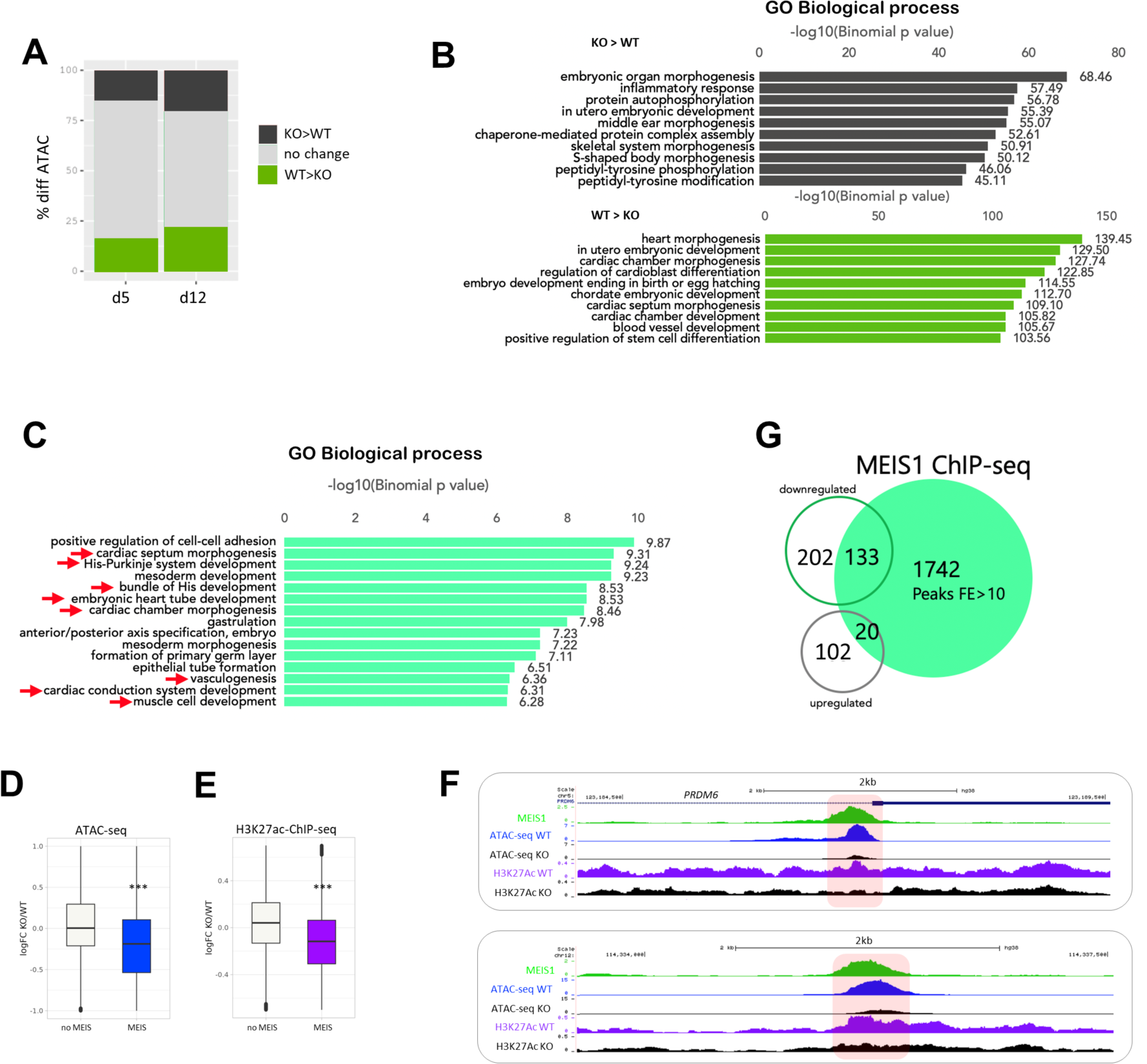
MEIS TFs promote chromatin accessibility and enhancer activation at critical regions for cardiac differentiation. A. Bar chart of chromatin accessibility shows unchanged accessibility at most chromatin regions in the absence of MEIS (d5 and d12). B. GREAT analysis of differential ATAC peaks (d5). Regions more accessible in wild-type (WT>KO, green bars) are associated to cardiac biological processes. In contrast ATAC-seq peaks higher in mutant (KO>WT, dark grey bars) shows association with genes involved in a variety of biological processes, including non-cardiac organ development and inflammation. The length of the bars corresponds to the binomial raw (uncorrected) P-values (x-axis values). C. GREAT analysis of MEIS1 high-confidence peaks (FE>10). MEIS1 peaks predominantly cluster around genes specifically associated with cardiac biological processes (red arrows). DE. Global changes in accessibility (D) and H3K27ac (E) in wild-type versus MEIS KO cells at d5 of cardiac differentiation, expressed as log2FC of normalized count values. The absence of MEIS TFs causes a significant reduction in accessibility and H3K27Ac at MEIS1 occupied regions, relative to unbound regions (One-sided t-test; *** = p<0.001). F. Loss of accessibility and acetylation at MEIS regulated genes. UCSC tracks of MEIS1 binding (green), ATAC-seq in wild-type (blue) and MEIS KO (black) and H3K27Ac ChIP-seq in wild-type (purple) and MEIS KO (black) at representative loci, *PRDM6* (top) and *TBX5* (*TBX5* transcription termination site is located 16kb downstream). Differential peaks are shaded in pink. G. MEIS1 predominantly function as an activator. Intersection of genes down- and up-regulated at d5 with genes associated to MEIS1 high-confidence peaks (using GREAT rules). The majority of genes dysregulated in MEIS KO and exhibiting MEIS1 binding nearby are activated by MEIS TFs (down-regulated in MEIS KO).

### MEIS cooperates with cardiac TFs

MEIS TFs are expressed in many non-cardiac cell types, yet they initiate a cardiac-specific transcriptional program. How do broadly expressed TFs execute lineage-specific functions? MEIS recognition motifs are enriched in tissue-specific enhancers, adjacent to motifs recognized by tissue-specific TFs (Fig. 1A). We hypothesized that MEIS could achieve tissue-specific functions through collaboration with lineage-specific partner TFs. To explore MEIS combinatorial binding in cardiac progenitors, we analyzed MEIS1 peaks using de novo motif discovery software [22]. We found MEIS recognition site as the second most enriched motif in MEIS1 peaks, with the top ranked motif corresponding to HAND2 motif. GATA, HOX-PBX and ZIC3 were also included in the top five overrepresented motifs (Fig 4A). Regions that lose accessibility in the absence of MEIS are enriched in largely similar motifs to those found in high confidence MEIS peaks in cardiac progenitors (Fig. S4A). Remarkably, regions that gain accessibility when transitioning from mesoderm to the cardiac progenitor state [15], are enriched in the same motifs (Fig. S4B), indicating that MEIS TFs operate as a hub in the network active in cardiac progenitors, which is fairly simple and dominated by few TFs at this stage. Next, we inspected up- and down-regulated genes at d5 and d7, to identify TFs whose expression is directly regulated by MEIS. *HOXB1, HOXB2, GATA4* and *GATA6*, whose recognition sites are enriched in MEIS1 ChIP-seq and in regions that gain accessibility in cardiac progenitors, are also activated by MEIS1-2, with HAND TFs being a notable exception (Fig S4C). Thus, MEIS positively control the expression of cardiac network TFs and cooperate with them.

**Figure 4.**
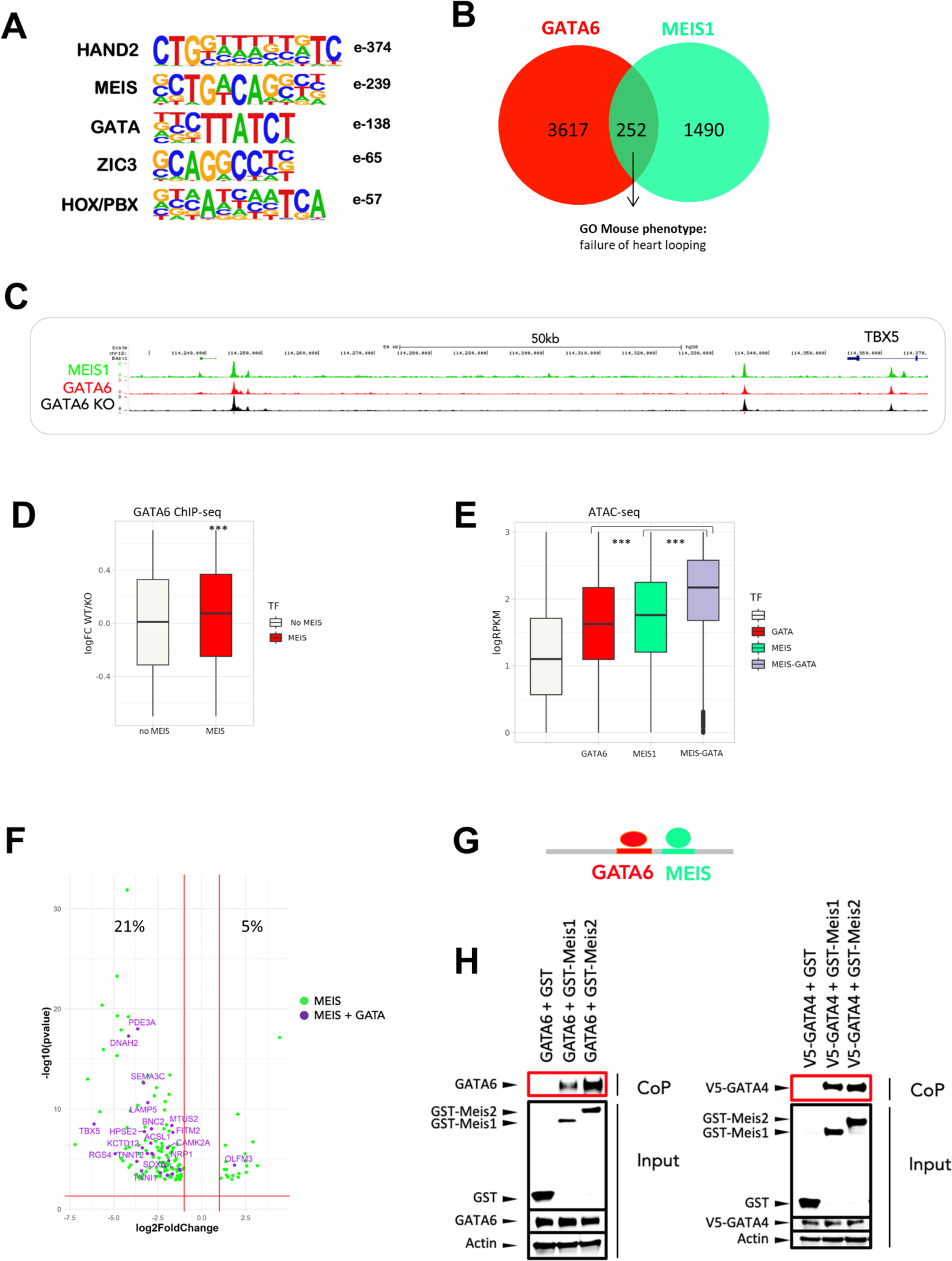
MEIS direct cooperativity with GATA TFs. A. Top motifs, ranked by significance, identified in MEIS1 peaks using de novo motif discovery (Homer). B. Non-proportional Venn diagram generated using MEIS1 and GATA6 top peaks (FE>10). MEIS1-GATA6 co-occupied regions are linked to genes whose loss of function in mouse leads to heart looping failure. C. UCSC tracks of MEIS1 (green) and GATA6 in wild-type and MEIS KO cells (red and black, respectively) at the *TBX5* locus. GATA6 peaks are still detected in the absence of MEIS1-2 (see also Fig S3). D. Changes in GATA6 binding levels at MEIS-bound location. A box plot of GATA6 differential binding (logFC wild-type vs. MEIS KO) shows that MEIS proteins increase GATA6 binding at MEIS1-occupied sites. In contrast, GATA6 binding at MEIS1-unbound regions is unaffected by MEIS proteins (One-sided t-test; *** = p<0.001). E. Co-binding of GATA6 and MEIS1 increases chromatin accessibility. A boxplot of accessible chromatin levels at GATA6 and MEIS1 occupied regions. Globally, regions co-occupied by GATA6 and MEIS1 have significantly higher accessibility relative to regions occupied by either MEIS1 or GATA6 (One-sided t-test; *** = p<0.001). F. Genes co-regulated by MEIS and GATA4 TFs in cardiac progenitors. Volcano plot of DE genes in MEIS KO that are also associated with a MEIS1 peak in d5 cardiac progenitors. Purple dots indicate genes that are also downregulated in GATA4 KO d6 iPSCs-derived cardiac progenitors; data sourced from [26]. Green dots are MEIS only targets. G. Model of indirect cooperativity: co-binding of GATA and MEIS TFs is guided by the vicinity of their recognition motifs. H. Co-precipitation assays. HEK293 cells were co-transfected with expression vectors for GATA6 or V5-tagged GATA4 and GST-tagged MEIS1 or MEIS2 or GST alone. Protein interactions were assayed by co-precipitation on glutathione beads directed toward the GST tag and eluted proteins analysed by western blotting to detect the presence of GATA6 and V5-GATA4 (red box, CoP). Cell lysates were analysed by western blotting prior to co-precipitation to detect protein expression (input), including ubiquitously expressed actin, used as a control.

To explore MEIS connectivity, we initially focused on GATA TFs. Given the established role of GATA4-5-6 in cardiac development [23], collaboration with these TFs could potentially explain how MEIS operate specifically at cardiac enhancers. Supporting MEIS-GATA combinatorial binding, GATA recognition motif is enriched in MEIS peaks (Fig 4A), and MEIS and GATA sites co-occur in cardiac developmental enhancers (Fig 1A). GATA motif is recognized by all six members of the GATA family [23], with *GATA4*, *GATA5*, and *GATA6* more highly expressed in d5 cardiac progenitors (Fig S4D). Experiments in mice and zebrafish suggest that the development of the cardiac system depends primarily on the overall dosage threshold of GATA proteins rather than on a specific GATA protein [24,25]. To investigate MEIS and GATA connectivity, we mapped GATA6 binding in d5 wild-type and MEIS KO cardiac progenitors. Next, we intersected high confidence MEIS1 and GATA6 peaks (FE>10) to establish if MEIS1 and GATA occupy a set of common regions. A limited fraction of top MEIS1 peaks (16%) overlaps GATA6 binding (Fig 4B; Table S2), yet these instances of co-binding are strategically positioned at critical genes for cardiac development and differentiation, including *TBX5, TNNT2, NRP1*, and also *HOXB1* (Fig 4C and S4E). Enhanced TF binding levels and chromatin accessibility are hallmarks of TF cooperativity because TFs that bind together reinforce each other’s occupancy on chromatin, outcompeting nucleosome for DNA access. We first examined if MEIS influences GATA6 chromatin occupancy. GATA6 binding signal is significantly higher in the presence of MEIS1-2 (Fig 4D). This increase is specifically observed when GATA6 occupies the same chromatin regions as MEIS1 and is therefore unlikely to be explained solely by the reduction in *GATA6* levels observed in MEIS KO. In contrast, the spatial distribution of GATA6 is largely unchanged in the absence of MEIS (Fig S4F, see also Figs 4C and S4E). Additionally, sites co-occupied by MEIS1 and GATA6 exhibit significantly higher accessibility compared to sites occupied by MEIS1 alone or GATA6 alone (Fig 4E). These findings collectively support cooperative binding interactions between MEIS and GATA. To address if MEIS-GATA combined binding has a functional impact, we analysed data from a GATA4 mutant model in iPSC-derived cardiac progenitors [26]. Notably, we found enrichment of MEIS motifs in GATA4 peaks within this system (Fig S4G). Consistent with functional synergy, 21% of MEIS direct target genes (downregulated in MEIS KO) exhibit downregulation in GATA4 loss-of-function within iPSC-derived cardiac progenitors (Fig 4F). Of note, the observation that GATA6 binds to genes co-regulated by MEIS and GATA4, even in the absence of MEIS (Fig 4C and S4E), suggests that GATA occupancy alone is not sufficient; MEIS is required for full gene activation. The enrichment of GATA motifs in MEIS peaks, and vice versa, implies a model wherein MEIS and GATA recruitment to shared enhancers is DNA-guided (Fig 4G). In addition, when overexpressed, both MEIS1 and MEIS2 can immunoprecipitate GATA4 and GATA6, indicating MEIS and GATA TFs can also form a complex (Fig 4H). In summary, the presence of nearby recognition motifs and protein-protein interactions underlie MEIS and GATA combinatorial binding, resulting in increased chromatin accessibility and the coordinated regulation of genes crucial for cardiac development and differentiation.

### Dynamic competition in MEIS combinatorial binding: balancing HOX and GATA interactions

MEIS peaks are also enriched in motifs recognized by HOX TFs and *HOXB1* and *HOXB2* are significantly downregulated in MEIS KO (Fig 4A; Fig S4C). The interaction between HOX and MEIS TFs is well established in a variety of organisms and tissues [8], but its significance in cardiac development remains underexplored. The human HOX family includes 39 members, which recognize highly similar sequence motifs [27]. We first asked which of the HOX genes are expressed in d5 cardiac progenitors. We found that anterior members of the HOXB cluster exhibit the highest expression levels (Fig S5A), including *HOXB1* and *HOXB2* that are significantly downregulated in the MEIS KO (Fig S4C). In mouse, overexpression of *HOXB1* in the second heart field (SHF) results in arrested cardiac differentiation; the effects of the overexpression are entirely consistent with the phenotype of *Hoxa1^-/-^; Hoxb1^-/-^* loss-of-function hearts [28]. To investigate the role of HOX proteins in this system, we chose a similar overexpression approach to circumvent the issue of redundancy often encountered in HOX mutants. We generated lines where *HOXB1* is placed under a doxycycline inducible promoter at a safe harbour locus [29]. *HOXB1* expression was induced just before the onset of *MEIS1* expression and maintained until d9 of cardiac differentiation (Fig 5A). At d9, elevated HOXB1 dosage led to a reduction in GFP-positive, NKX2-5 expressing cells, which exhibited little to no contraction compared to their control counterparts, indicating a delay in cardiac differentiation. In addition to *NKX2-5*, the downregulation of key genes like *TNNT2* and *MYH6*, encoding for cardiac-specific troponin and myosin, which function in cardiac muscle contraction, further underscores HOXB1 inhibitory effect on cardiac muscle differentiation (Fig 5B). In sum, overexpression of *HOXB1* has a negative effect on cardiac differentiation, which is consistent with *Hoxb1* overexpression in vivo and in cardiac differentiation of mouse ESC [28]. Unsurprisingly, both MEIS1 and MEIS2 demonstrate the capacity to interact with HOXB1, given their well-established partnership with HOX proteins (Fig 5C). Cardiomyocyte differentiation involves several regulatory waves that lead to activation of progenitor-specific genes followed by activation of cardiac-specific genes. We examined *HOXB1* and *GATA* expression in a time course of cardiac differentiation. We found distinct expression dynamics: *HOXB1* expression peaks during the progenitor stage (d5 - d7) and then declines to basal levels in cardiomyocytes. Conversely, expression of *GATA4* increases with cardiomyocyte differentiation, and both *GATA4* and *GATA6* remain highly expressed in cardiomyocytes (Fig 5D). These contrasting expression dynamics suggest that HOX and GATA may act independently to control distinct aspects of cardiac lineage differentiation, rather than acting in concert. Consistent with this hypothesis, HOXB1 and GATA exhibit seemingly opposite effects on cardiac differentiation: while HOXB1 delays cardiac differentiation, GATA4-5-6 are known for their established cardiogenic effects, with GATA4 being one of three TFs capable of converting fibroblasts into cardiomyocytes [30]. These observations, together with the finding that GATA4 and HOXB1 interact with MEIS1 (Fig 4H and Fig 5C), suggest a potential competition for MEIS binding between these two TF families. Supporting this hypothesis, the addition of HOXB1 reduced the level of GATA4 co-precipitated by MEIS1 to 40% (Fig 5E). Similarly, the addition of GATA4 decreased the amount of HOXB1 co-precipitated by MEIS1. To test if HOX-GATA competitive binding affects MEIS genomic occupancy, we altered HOX-GATA balance during cardiac differentiation. We used our HOXB1 overexpression model to maintain high HOXB1 dosage up to d8, when HOXB1 levels are declining and GATA4 expression is steadily increasing. We first selected HOXB1-MEIS target enhancers based on specific criteria, including overlap with a top MEIS1 peak, increased accessibility in wild-type cells, presence of at least one HOX-PBX consensus motif, and association with a DE gene in MEIS KO cells. Subsequently, we validated HOXB1 binding to these regions using ChIP-seq (Fig 5F). Next, we identified GATA-MEIS target enhancers, defined as regions bound by GATA4 and GATA6 (Fig 5F), overlapping with MEIS1 top peaks and linked to genes downregulated in GATA4 KO cardiac progenitors [26]. We found that sustained high dosage of HOXB1 in d8 cardiomyocytes leads to increased MEIS binding levels at HOXB1-bound enhancers (Fig 5G; Fig S5B). The expression of genes associated with these enhancers is also increased, suggesting a functional effect (Fig 5H). Conversely, in the same HOXB1 overexpressing cells, we observed a reduction in MEIS1 binding levels at high-confidence GATA enhancers, correlating with decreased expression of genes associated with these enhancers (Fig 5GH; Fig S5B). These results suggest that HOXB1 may delay cardiac differentiation by diverting MEIS away from GATA-bound, cardiac-specific enhancers. Supporting this interpretation, most HOX-MEIS binding is excluded from GATA-MEIS regions and is associated with regulation of various processes, not just those related to cardiac differentiation, as observed for GATA-MEIS regions (Fig S5C). In sum, MEIS acts as a molecular switch, trading partners to control the progression of cardiac differentiation. A similar separation of GATA6-MEIS and HOXA3-MEIS binding, each associated with distinct biological processes, is observed in the cardiac neural crest-populated posterior branchial arches (PBA) [31] (Fig S5D). This indicates that HOX and GATA diversify MEIS’s regulatory functions and suggests that their mutual competition for MEIS is a general mechanism.

**Figure 5.**
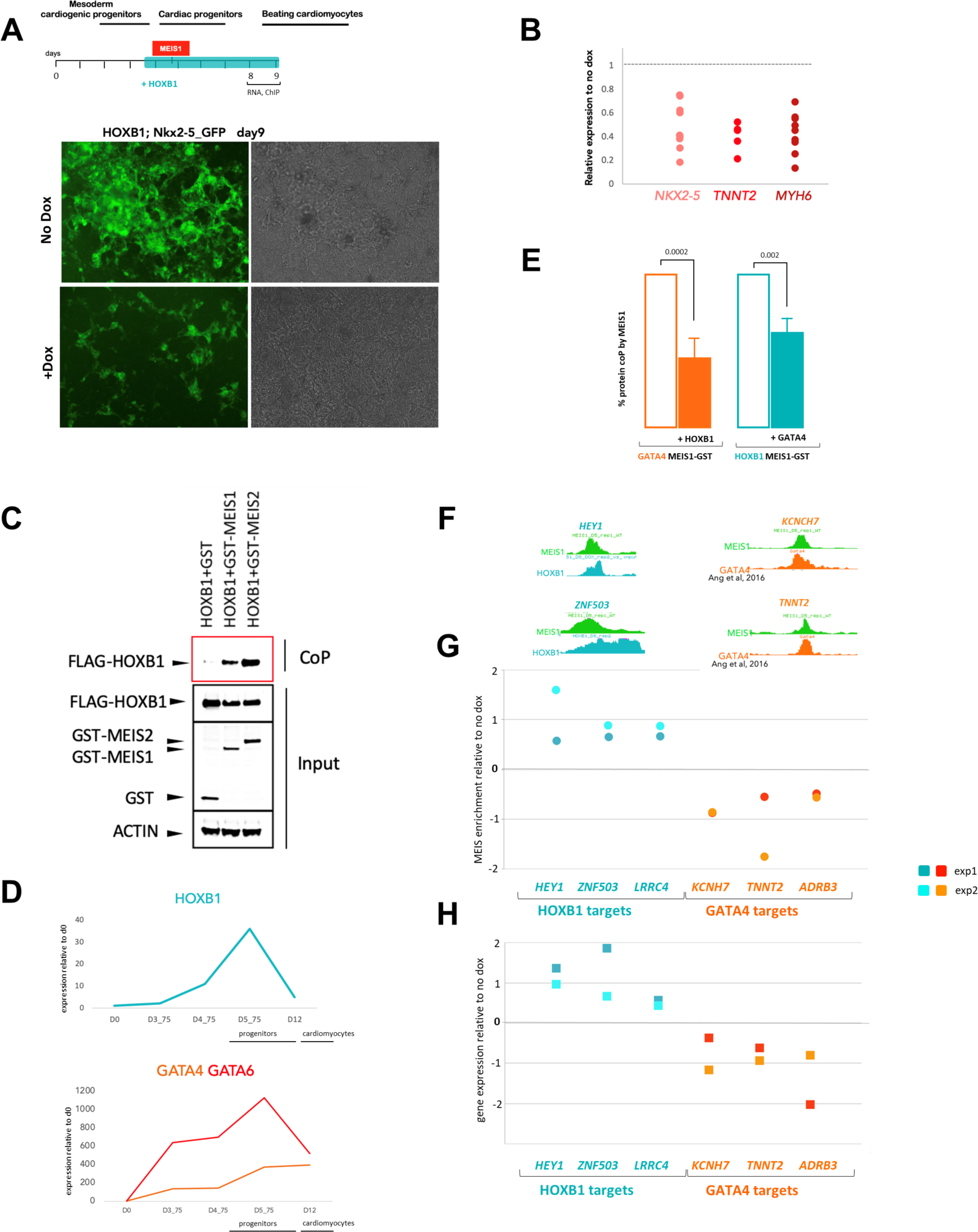
MEIS combinatorial binding drives the dynamics of cardiac lineage differentiation. A. Schematic of *HOXB1* overexpression along the timeline of cardiac differentiation. At d9, HOXB1; NKX2-5-GFP cells display strong GFP expression, mirroring NKX2-5 expression. HOXB1 induction with doxycycline results in fewer cells expressing GFP. B. Cardiomyocyte markers are downregulated in d9 HOXB1-overexpressing cells. *NKX2-5, MYH6* and *TNNT2* expression, measured by qPCR, is expressed as a percentage relative to their expression measured in control, non-induced cells. C. Co-precipitation assays. HEK293 cells were co-transfected with expression vectors for FLAG-tagged HOXB1 and GST-tagged MEIS1 or MEIS2, or GST alone. Protein interactions were assayed by co-precipitation on glutathione beads directed toward the GST tag and eluted proteins analysed by western blotting to detect the presence of FLAG-HOXB1 (red box, CoP). Cell lysates were analysed by western blotting prior to co-precipitation to detect protein expression (input), including ubiquitously expressed actin, used as a control. D. Dynamic expression of *HOXB1* and *GATA4-6* during cardiac differentiation. Time course of cardiac differentiation analysed by RNA-seq. At each timepoint shown, normalised transcript counts are expressed as fold change relative to their value at the start of differentiation (d0). E. HEK293 cells were co-transfected with expression vectors for GATA4, GST-tagged MEIS1, and FLAG-tagged HOXB1. Co-precipitation of GATA4 with MEIS1-GST in the presence of HOXB1-FLAG was detected by western blotting and is expressed as a relative percentage of GATA4 co-precipitated with MEIS1-GST in the absence of HOXB1 (orange bars; Tukey’s test p < 0.0002). Co-precipitation of HOXB1 with MEIS1-GST in the presence of GATA4 was detected by western blotting and is expressed as a relative percentage of HOXB1 co-precipitated with MEIS1-GST in the absence of GATA4 (teal bars; Tukey’s test p < 0.002). F. UCSC tracks of MEIS1 (green) in d5 cardiac progenitors, HOXB1 FLAG (teal) in d8 cardiomyocytes and GATA4 (orange) in d6 cardiac progenitors at representative HOXB1-MEIS1 target regions (left) and GATA-MEIS regions (right). GH. MEIS1 binding levels (G) and gene expression (H) in d8 cardiomyocytes with sustained HOXB1 dosage. G. ChIP qPCR using MEIS1 antibody in d8 dox-induced (HOXB1 overexpressing) and -dox (control) differentiating cardiomyocytes. Scatter plot of MEIS1 binding levels measured at HOX-MEIS target regions (blue) and GATA-MEIS bound regions (orange). MEIS1 enrichment (relative to input) is plotted as log FC relative to MEIS1 enrichment measured at the same region in the control. Blue dots represent log FC value of MEIS1 enrichment measured on HOXB1-target enhancers in two independent experiments. Orange and red dots represent log FC value of MEIS1, measured on GATA4-target enhancers in two independent experiments; see also Fig S5B). H. Expression of *HEY1, ZNF503, LRRC4* (MEIS1-HOXB1 targets) and *KNCH7, TNNT2, ADRB3* (MEIS1-GATA4 targets), measured by qPCR, and expressed as log FC of their relative expression measured in control, non-induced cells.

### MEIS promotes accumulation of KMT2D at cardiac enhancers

While it is well established that lineage-specific TFs are essential for lineage-specific transcription, our results indicate that broadly expressed TFs are also essential. What function do they provide in cell differentiation programs? Based on the association of MEIS1 binding with transcriptional activation, a likely hypothesis is that MEIS TFs harness a general function, critical for the activation of gene expression. A clue to the nature of this general function comes from the observation that MEIS2 syndrome, caused by mutations in the *MEIS2* gene, display clinical features – distinctive facial appearance, developmental delay, intellectual disability, heart and skeletal abnormalities - that closely resembles those of Kabuki syndrome [32]. Kabuki syndrome is caused by mutation in the *KMT2D* gene [33], encoding for a ubiquitously expressed histone-lysine N-methyltransferase, which monomethylates lysine 4 of histone H3 (H3K4me1) of distal enhancers, leading to transcriptional activation [34]. KMT2D lacks a DNA binding domain and relies on DNA binding proteins to identify its correct genomic addresses [35]. Similar to MEIS, KMT2D is required for cardiac development [36]. We surveyed available datasets for evidence of an interaction between MEIS1 and KMT2D. Dysregulation of KMT2 proteins (MLL) and MEIS1 contribute to acute myeloid leukemia (AML). A physical association between MEIS1 and KMT2D (MLL4) has been reported in AML cell lines [37], where KMT2D is identified among the top 50 MEIS1 interactors. Next, we asked if MEIS1 and KMT2D bind common regions. We observe high-confidence KMT2D ChIP-seq signal at top MEIS1-bound regions in cardiac progenitors, suggesting coordinate binding between MEIS1 and KMT2D. KMT2D peaks associated with MEIS1 binding are found nearby genes involved in cardiac development and differentiation, in contrast to the remaining KMT2D peaks, mostly associated with metabolic pathways (Fig 6A). We confirmed MEIS1 chromatin colocalization with KMT2D using reChIP (Fig 6B). These results indicate that MEIS1 and KMT2D bind together to a subset of regions. To establish causality, we interrogated KMT2D binding in the absence of MEIS1-2. We found that KMT2D levels are significantly higher in wild-type cells relative to MEIS KO cells, and this effect is specific to regions bound by MEIS1 (Fig 6C). This trend is exemplified by the *ZNF503* locus, where KMT2D signal is specifically decreased at regions overlapping MEIS1 peak. In contrast KMT2D binding at regions not overlapping MEIS1 peaks appears unchanged (Fig 6D). Together, these observations suggest that MEIS1-2 function to recruit KMT2D at lineage-specific enhancers to activate the cardiac gene expression program. KMT2D monomethylates histone H3 at enhancers active during cell fate transitions, eventually promoting recruitment of p300 and deposition of H3K27Ac [38,39]. We explored the distribution of H3K27Ac signals in wild type cells relative to MEIS KO. H327Ac signal is significantly higher at regions where both MEIS1 and KMT2D are present indicating a synergy between MEIS1 and KMT2D to enhance chromatin acetylation (Fig 6E). In summary, MEIS proteins function as *actuator* TFs in cardiac cells. They initiate cardiac-specific transcription by harnessing general co-activators at cardiac-specific enhancers, to which they directed by binding with lineage- and/or domain-specific TFs (Fig 7).

**Figure 6.**
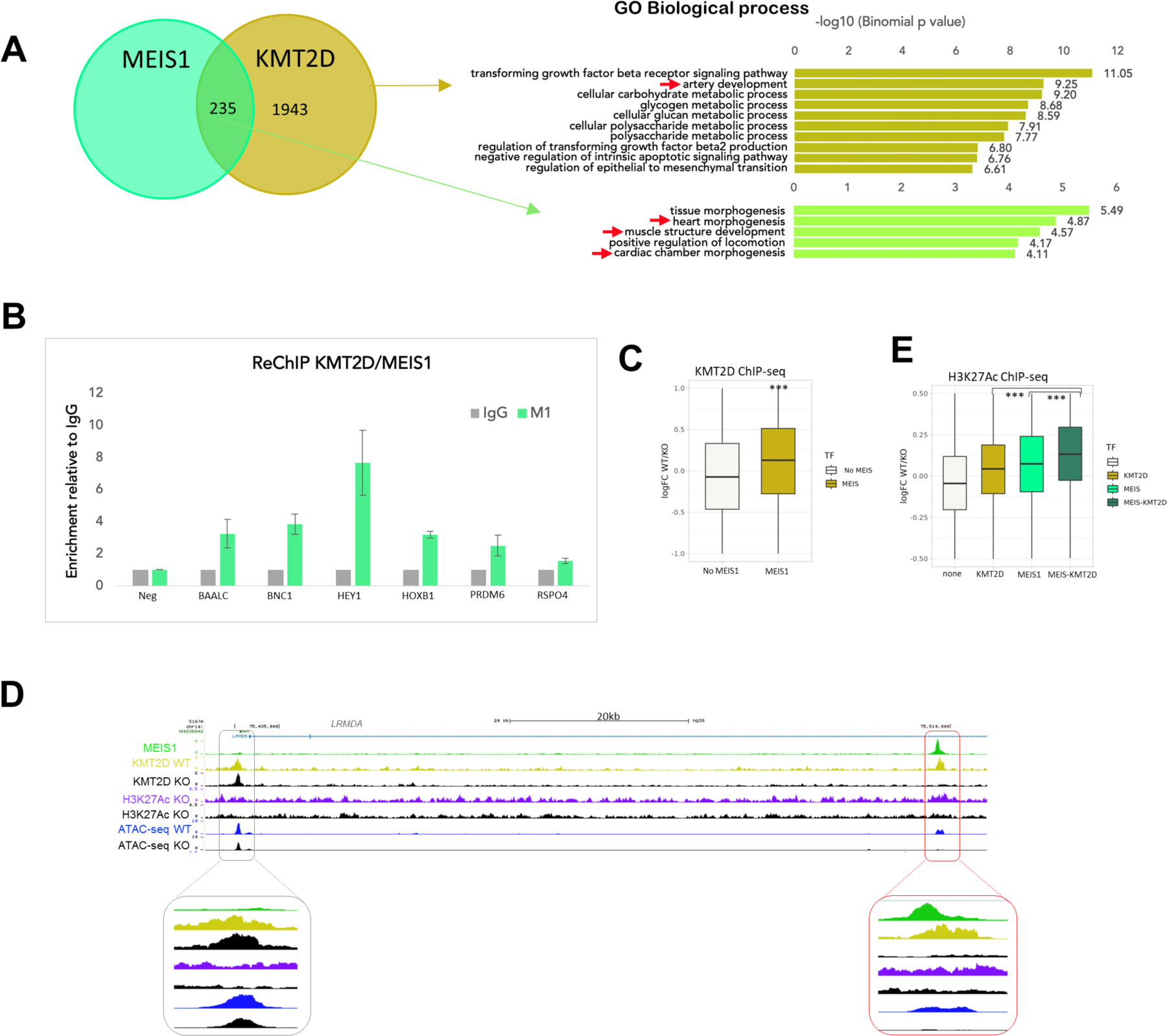
MEIS promotes accumulation of KMT2D at cardiac enhancers. A. Intersection of MEIS1 and KMT2D top peaks (FE>10). Shared MEIS1-KMT2D peaks are mainly associated with genes implicated in cardiac GO. In contrast, KMT2D only peaks cluster around genes involved in metabolic processes. B. Re-ChIP using KMT2D antibody, followed by MEIS1 antibody. Chromatin obtained from KMT2D immunoprecipitated chromatin was again precipitated using MEIS1 or an unspecific (IgG) antibody and amplified using primers spanning KMT2D and MEIS1 co-occupied regions. KMT2D-occupied regions are enriched upon MEIS1 immunoprecipitation, while no enrichment is observed using a negative control region; the negative control is an unbound region. MEIS1 binding enrichment is expressed relative to IgG. C. Changes in KMT2D occupancy in MEIS KO cells. Boxplot of KMT2D logFC in wildtype versus KO cells. The presence of MEIS TFs significantly increases KMT2D signal at MEIS1-occupied regions. In contrast, no significant change is observed at regions that are not bound by MEIS1 (One-sided t-test; *** = p<0.001). D. UCSC tracks of MEIS1 binding (green), ATAC-seq in wild-type (blue) and MEIS KO (black), H3K27Ac ChIP-seq in wild-type (purple) and MEIS KO (black) and KMT2D ChIP-seq in wild-type (gold) and MEIS KO (black) at the *ZNF503* locus. Differential KMT2D peaks, boxed in red, overlap MEIS1 peaks. In contrast, loss of MEIS1 does not affect KMT2D binding at MEIS1 unbound regions (boxed in grey). Enlarged views of the same regions are shown below the tracks. E. Boxplot of H327Ac logFC in wildtype versus KO cells. Relative to regions occupied by KMT2D or MEIS1 alone, regions co-occupied by MEIS1 and KMT2D display increased acetylation in wildtype versus KO cells, suggesting that co-occupancy of MEIS1 and KMT2D enhances chromatin acetylation (One-sided t-test; *** = p<0.001).

**Fig 7.**
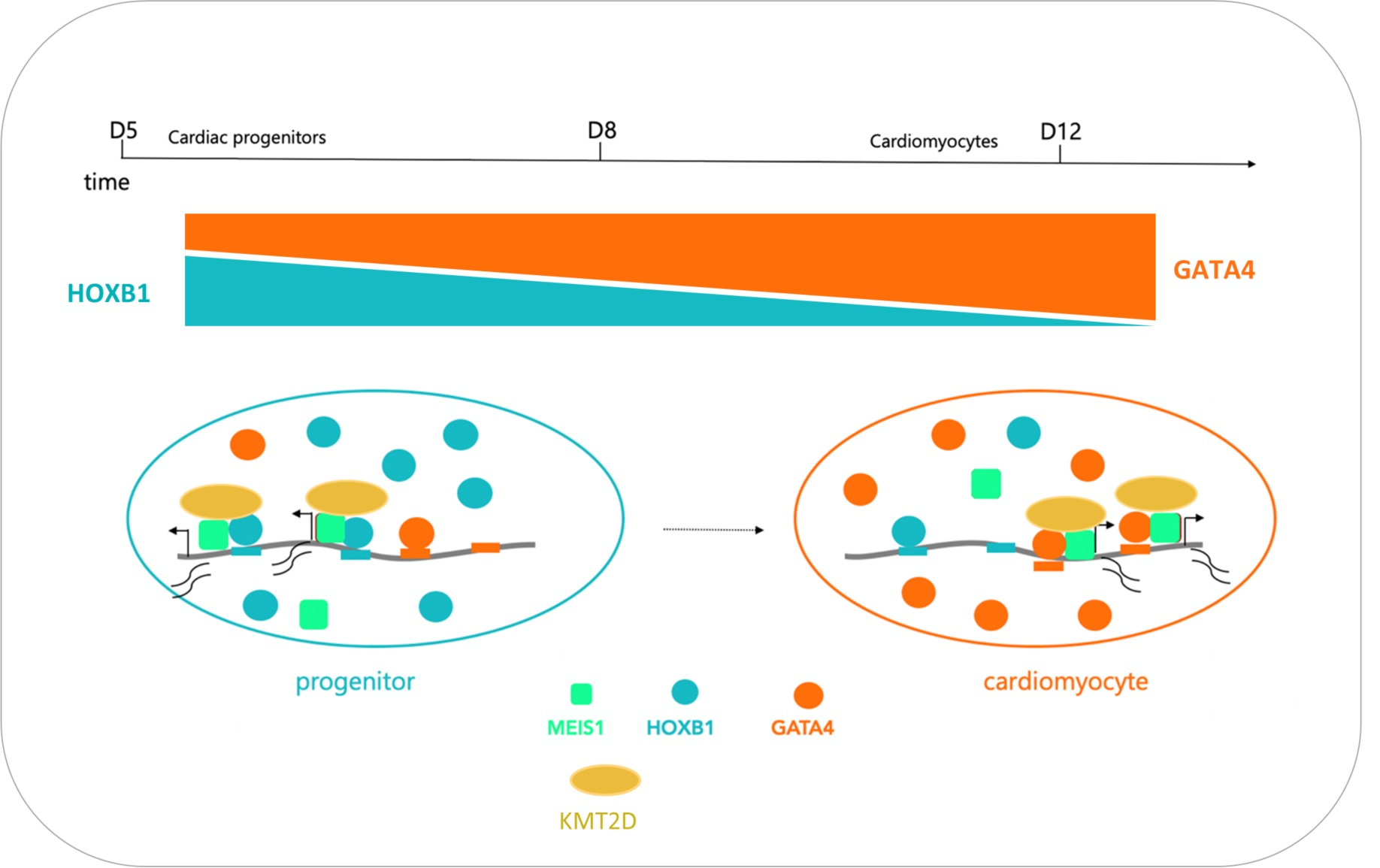
Model. Variation in TF dosage regulate the distribution of MEIS actuators at enhancers. At progenitor stage, HOXB1 levels are high, GATA4 are low. MEIS interacts predominantly with HOX, which directs MEIS to its target enhancers. Here MEIS1 promotes accumulation of KMT2D leading to activation of HOX target genes. As differentiation proceeds, GATA4 levels increase and HOXB1 levels decrease. This favours MEIS interaction with GATA; GATA4 directs MEIS to function at its target enhancers to drive expression of its target genes. Combinatorial binding is dynamic and associated with the progression of cardiac differentiation: MEIS actuators fully activate an ‘early’ cardiac progenitor program in concert with HOX and a subsequent specification module together with GATA TFs.

## Discussion

Combinatorial binding is a common feature of mammalian transcription networks, where TFs collaborate to access DNA and enhance their chromatin binding. This process allows for binding and response specificity, broadening the regulatory capabilities of the finite set of TFs encoded by an organism. However, it is less clear if combinatorial binding also functions to assemble complexes with emerging functional properties in transcriptional activation. Here, we identify the essential role of the broadly expressed, non-lineage-specific TFs MEIS in activating lineage-specific transcription and establishing cardiac fate. We propose that MEIS proteins are obligatory components of productive TF combinatorial binding, functioning as actuator TFs to effectively activate the transcriptional programs selected by lineage- or domain-enriched TFs (GATA and HOX). MEIS TFs, though largely ubiquitous, are selectively guided to cardiac-specific enhancers by binding with lineage-enriched TFs. Increasing the binding levels of actuator TFs at cardiac enhancers activates gene expression. This happens, at least in part, because MEIS TFs promote accumulation of the methyltransferase KMT2D to initiate enhancer commissioning.

MEIS TFs significantly promote transcription, predominantly eliciting positive effects. This aligns with previous observations of high MEIS binding correlation with enhancer activity and gene activation across different tissues of the developing mouse embryo [10]. TALE TFs are evolutionarily ancient and were present in unicellular eukaryotes before the diversification of metazoans [40,41]. An intriguing possibility is that, with the advent of multicellularity and the generation of specialized cell types, new tissue-restricted proteins (e.g. HOX) evolved within the genetic toolkit of animals to harness the function of these ancient activators in a cell type- or tissue-specific manner.

This new framework suggests a strategy for efficient cell reprogramming. Lineage-specific TFs are the primary candidates for stem cell differentiation and reprogramming protocols, yet combinations of these TFs or those enriched in specific lineages often prove inefficient in generating the desired cell type [2]. Combining lineage-specific TFs with broadly expressed TFs, which are often overlooked in tissue-specific processes, could significantly enhance the efficiency of direct reprogramming.

Congenital heart disease represents a significant birth defect, impacting up to 1% of all live births in the Western world [42]. Extensive research into cardiac development and differentiation has identified numerous TFs and signalling pathways as critical components in vertebrate heart formation [43]. Previous studies have suggested a potential role for MEIS proteins in regulating cardiomyocyte differentiation [44-48], but the requirement of MEIS TFs for human cardiac differentiation has remained largely unrecognized. Here, we identify MEIS TFs as essential regulators of human cardiomyocyte differentiation. MEIS function as a central hub, enabling and coordinating the progression of cardiac differentiation.

MEIS KO has a profound impact on cardiac differentiation, affecting both cardiac and epicardial lineages. Motif analysis indicates that MEIS co-binding partners are also involved in the transition from mesoderm to cardiac progenitors, highlighting MEIS as a key node of cardiac transcription networks. MEIS1 and MEIS2 can physically associate with HOXB1 and GATA4-6. GATA and HOX recognition motifs are enriched in MEIS peaks; no evidence of combined binding sites was found, indicating that complex formation does not alter the binding capabilities of its individual components. Several lines of evidence suggest that MEIS can bind alternately, but not simultaneously, with HOX and GATA. Firstly, GATA and HOX can outcompete each other for binding to MEIS. Secondly, HOX and GATA exhibit opposite effects on cardiac differentiation. Thirdly, a dynamic temporal expression pattern is observed: HOX levels are high in cardiac progenitors and low in cardiomyocytes, while GATA4 levels show the opposite pattern. Variations in TF dosage regulate the distribution of MEIS at enhancers. Maintaining high levels of HOX beyond the progenitor stage, delays cardiac differentiation. At the genomic level, MEIS is redistributed from GATA-bound enhancers to HOX-bound enhancers, leading to changes in the expression of the associated genes. HOX proteins are well-established partners of MEIS proteins [8]. They control embryonic patterning in all animals, deploying different mechanisms, including control of cell fate [27]. In mouse, HOXB1 maintains cardiac progenitors, as its depletion accelerates cardiac differentiation [28]. Given the central role of MEIS in cardiac differentiation, one possible mechanism for HOX action is to restrict the availability of MEIS, preventing it from interacting with cardiac TFs like GATA. This restriction may help to maintain cells in a progenitor state, allowing for sufficient proliferation before committed progenitors enter terminal differentiation. Therefore, we propose that the dynamics of MEIS combinatorial binding, influenced by changes in the dosage of its binding partners, orchestrate the progression of cardiac differentiation.

A number of observations suggest that MEIS control the cardiac differentiation transcriptional program in concert with TBX5, in addition to GATA TFs. MEIS1 is among the top TBX5 interactors in day 6 cardiac progenitors [26]. TBX TFs recognise a sequence (TGNTGACA) which largely coincides with TALE motif (TGACA), and MEIS and TBX5 co-occupancy has been established in the limb [49]. Finally, TBX5 and GATA4 show extensive binding overlap, and functionally interact to drive cardiac morphogenesis [26,50,51]. All of the above suggests that MEIS cooperate with the key TFs active in cardiac progenitors GATA4-6, and TBX5.

MEIS TFs are widely expressed and, beyond cardiac differentiation, are indispensable for the development of multiple organs and can exhibit oncogenic behaviour. MEIS cooperate with tissue-restricted TFs in different contexts. For instance, in the branchial arches, the homeodomain TF HOXA2 activates its transcriptional program by enhancing MEIS TF binding to potentially lower-affinity sites throughout the genome [52]. In projection neurons, MEIS2 is guided to enhancers associated with projection-neuron-specific genes by the homeodomain TF DLX5 [53]. In the retina, MEIS TFs collaborate with the homeodomain TF LHX2 to activate the expression of retinal progenitor cell-specific genes [54]. MEIS TFs cooperate with HOX and TBX5 to establish early limb development [49]. These examples support the model that tissue-specific transcriptional activation results from the combined effect of ubiquitously expressed actuator TFs and lineage- and/or domain-specific TFs.

In summary, our results reveal that that combinatorial binding assembles widely expressed actuators and lineage-restricted TFs into selective, transcriptionally active complexes. Tissue-specific transcription results from ubiquitous actuators harnessing general transcriptional activators at tissue-specific enhancers, to which they directed by binding with lineage- and/or domain-specific TFs. This model offers insights into how TF synergy shapes transcriptional regulation.

## Acknowledgements

We thank Andy Hayes, Leo Zeef and members of the Genomic Technologies, Bioinformatics and Bioimaging Core Facilities at the University of Manchester. We also thank Charles Sagerstrom and Andy Sharrocks and for feedback on the manuscript. This work was funded by the Biotechnology and Biological Sciences Research Council (BB/T007761 to NB and BB/X016684 to NB and MJB) and the UKRI Future Leaders Fellowship (MR/T041668/1 to MJB).

## Methods

### Maintenance of human ESC lines

HES3 *NKX2-5^eGFP/w^* hESCs [14] and gene-edited derivatives were maintained on a layer of mitotically-inactivated mouse embryonic fibroblasts (MEFs) in DMEM/F12-based medium containing 20% (v/v) KnockOut™ Serum Replacement (Gibco), 100 mM non-essential amino acids (Gibco), 2 mM GlutaMAX™ (Gibco), 0.1 mM β-mercaptoethanol (Gibco) and 10 ng/ml bFGF (Miltenyi Biotech). hPSCs were passaged using TrypLE Select (Thermo Fisher Scientific) every 3-4 days.

### hESC differentiation

Differentiations were performed in serum-free BPEL medium containing 1 μg/mL insulin using an embryoid body (EB) or monolayer format. 24 hours before differentiation was induced, hESCs were plated at a density of 7x10^4 / cm^2^ (for EBs) or 2x10^4 / cm^2^ (for monolayer) in normal growth medium in 6-well plates. EBs were then formed by depositing 2500 cells in 50 μl differentiation medium per well in V-bottomed 96-well plates (Greiner). The following growth factors were present for the first 3 days of differentiation: 25 ng/ml BMP4 (R&D Systems), 25 ng/mL Activin A (Miltenyi Biotech) and 1.5-1.75 μM CHIR99021 (Selleckchem). On day 3, wells were refreshed with 100 μl BPEL. On day 6, wells were refreshed with 100 μL BPEL supplemented with 100 pg/ml bFGF (Miltenyi Biotech). On day 9, wells were refreshed with 100 μL BPEL. Monolayers followed the same timing but were cultured in 6-well plates in 3.5 ml of medium and 5 µM XAV939 (Tocris) was included between day 3 and 6.

### Generation of MEIS1-2 knockout hESCs

Ribonucleoprotein (RNP) complexes of CRISPR-Cas9 were made by combining 120 pmol of crRNA:tracrRNA mix (IDT) with 67 pmol of Cas9 (Alt-R™ S.p. Cas9 Nuclease V3, IDT), final concentrations 1.3 and 1.1 µM respectively. gRNA sequences were *MEIS1*: ACGACGATCTACCCCATTAC and *MEIS2*: CTTCAAGGCGTCGTTGACAG. *MEIS1* and *MEIS2* were targeted sequentially. 0.5x10^6^ cells were transfected with the RNP mix and 120 pmol Alt-R® Cas9 Electroporation Enhancer (1.3 µM final). Transfection was performed by electroporation using an Amaxa Nucleofector II combined with Human Stem Cell Nucleofector Solution 2 (Lonza, ref #: VPH-5022) and programme B-16. 3 days later, single cells were sorted by FACS (BD Influx) into MEF-coated 96-well plates. DNA was isolated from clones using a Qiagen DNA Mini kit for PCR screening using primers spanning the cut sites. PCRs were performed using Q5 High-Fidelity DNA polymerase (NEB). Homozygous mutant clones were confirmed by Sanger sequencing using primers listed in Table S3.

### Generation of Tet-On-HOXB1 hESC line

Human embryonic stem cells NKX2-5eGFP/w (Elliott et al., 2011) were genetically modified by site-specific placement of Dox-inducible HOXB1 at the H11 locus using DICE system [29]. DICE system was introduced into the *H11* locus of NKX2-5eGFP/w using 2 plasmids: one carrying Cas9 nuclease and two guide RNAs (Addgene#164850) and a second one carrying a landing pad (Addgene #51546) with homology arms (HAs) for homologous recombination at the *H11* locus (Addgene #164851). The donor plasmid (Addgene #51547) carried *HOXB1* gene (GenScript NM_002144.4) under control of TET-ON promoter (Addgene 72835) and modified rtTA (Tet_On 3G) under control of the UbC promoter (Addgene#19780). Transfections were performed in 6-well plates overnight, using Lipofectamine™ Stem Transfection Reagent (ThermoFisher), following manufacturer instructions. For the generation of Pad_H11 cell line, 2mg landing pad plasmid and 1mg Cas9-sgRNA-encoding plasmid were used for Cas9-mediated recombination. For HOXB1 integration, 1mg was used for each of phiC31 integrase (Addgene #51553), Bxb1 integrase (Addgene#51552) and HOXB1 donor plasmids. Cells were selected with either 100 mg /ml G418 (Invitrogen) for HR or 1mg/ml puromycin (Invitrogen) for DICE, starting 2 days after transfection and selecting cells for 4 days. Clones were isolated and selected for their ability to efficiently upregulate HOXB1 on exposure to 1 μg/ml doxycycline (Sigma-Aldrich). Genetic modification was confirmed by genotyping, followed by sequencing using primers listed in Table S3.

### Real time RT-qPCR

Total RNA was isolated using a RNeasy kit (Qiagen) according to the manufacturer’s instructions and transcripts detected in a one-step RT–qPCR reaction using 30 ng of total RNA and Quantitect SYBR green reagent (Qiagen). Data were normalized against the control gene RPLPO or TUBB5. RNA samples were run in duplicate from at least three independent experiments. RT-PCR primers are listed in Table S3.

### ChIP and reChIP

MEIS1 ChIP was performed in HOXB1 overexpressing cells on d8 of cardiac differentiation as described previously [55] using MEIS1 antibody (Abcam ab 19867) and IgG Rabbit Polyclonal (Merck Millipore). Bound regions were detected by qPCR using the primers listed in Table S3.

For re-ChIP assay, we used 24x10^6^ NKX2-5eGFP/w cells, collected on d5 of cardiac differentiation. After the first round of immunoprecipitation with rabbit polyclonal anti KMT2D (Atlas Antibodies), the beads were washed and then incubated as described in [55] with minor modifications: for sequential ChIPs precipitated complexes were eluted twice in 55 ml Elution buffer (1xTE,2% SDS, 10mM DTT) containing fresh protease inhibitor cocktails for 30 min at 37 ℃ with shaking. 10% sample was set aside as total in-line control ChIPs. To decrease SDS concentration, volume was increased 20 times with ChIP buffer (1 mM EDTA, 0.5 mM EGTA, 4 mM Tris-HCl, pH8.0, 100 mM NaCl, 0.1% Na-deoxycholate, 17 mM N-lauryl sarcosine) and subjected to a second round of immunoprecipitation with MEIS1 antibodies or IgG control. Following reverse cross-linking and DNA purification, bound regions were detected by quantitative PCR (qPCR), using Quantitect SYBR green PCR reagent (Qiagen) and primers listed in Table S3. Results were analyzed with Rotorgene Software 2.3.5 (Corbett Research) and expressed as fold enrichment relative to IgG. Two independents experiments were performed.

### Coprecipitation experiments

Coding sequences for GATA4, GATA6, HOXB1, MEIS1b, MEIS2.1 were cloned into mammalian expression vectors. GATA4 has a V5 tag, HOBX1 has a FLAG tag, and MEIS1b and MEIS2.1 have a GST tag. Cells were seeded in 6-well plates at 400,000 cells/well and transfected after 24 hours using 1μg of plasmid and Fugene6 (#E2691, Promega) according to the manufacturer’s instructions. For experiments using GATA4, HOXB1, and MEIS1b, an empty pcDNA3 vector was used to keep DNA levels equal in each condition.

Proteins were collected in ice-cold IPLS lysis buffer including protease inhibitor cocktail (#11873580001, Roche) 48 hours after transfection. Glutathione-sepharose beads (#GE17-0756-01, Sigma) were pre-washed three times with ice-cold IPLS lysis buffer, and then incubated overnight with samples on a rotating wheel at 4 °C. Beads were washed three times with ice-cold IPLS. Input and CoP samples were boiled in Laemmli buffer, run on SDS-PAGE and visualized using anti-FLAG (M2) (#F1804, Sigma), anti-GATA4 (#19530-1-AP, Proteintech), anti-GATA6 (#5851, Cell Signalling), anti-GST (G1160, Sigma), HRP-conjugated anti-β-ACTIN (#A3854, Sigma), and HRP-conjugated secondary antibodies.

### Immunocytochemistry

EBs were fixed with 10% neutral buffered formalin (Merck) for 20 minutes, followed by 3X PBS washes. EBs were permeabilised with 0.5% Triton X-100/PBS for 1 hour and blocked for 1 hour with 4% donkey serum/PBS (Merck), both at room temperature. Primary antibodies against α-actinin (1:800; Merck A7811) and WT1 (1:200; Abcam ab89901) were incubated at 4°C in blocking buffer overnight. EBs were washed 3X with 0.05% PBS/Tween-20, followed by incubation with AlexaFluor secondary antibodies for 1 hour at room temperature. EBs were subsequently washed 3X with PBS/Tween-20 and nuclei stained with 1 µg/ml Hoechst for 15 minutes at room temperature. EBs were washed with PBS/0.1% BSA and mounted onto coverslips using ProLong™ Gold Antifade Mountant (Thermo Fisher Scientific). Images were acquired using SP8 inverted confocal microscope (Leica).

### Identification of TALE motif in developmental enhancers

TF motifs co-occurring with tissue-specific TF’s motifs at tissue-specific developmental enhancers were identified using the method described in Garcia-Mora et al. [12]. Tissue-specific TFs were identified using human embryonic H3K27ac ChIP-seq data and RNA-seq data [56,57]. Motifs corresponding to tissue-specific TFs were obtained from the Homer motif database under the names Gata4 (GATA_GATA4), Hand2 (BHLH_HAND1), Tbx20 (TBOX_TBX20), Olig2 (BHLH_MYOD1), Hoxc9 (HOMEOBOX_Hoxc9), RFX (RFX_RFX4),

Atoh1 (BHLH_ATOH1), Nr5a2 (NR_NR5A2). Coordinates of tissue-specific motifs in the human genome (hg38) were identified using scanMotifGenomeWide.pl script, from HOMER (v4.11, 10-24-2019) [22]. Using BEDtools [58], tissue-specific motif coordinates were identified in putative tissue-specific enhancers [57] to use as foreground. Tissue-specific motif coordinates found in random genomic regions were used as background for motif enrichment analysis. Motif enrichment analysis was performed using the findMotifGenomeWide.pl script, from HOMER using a size parameter of 200 nt. A p-value cut-off of 1x10^-5^ was used and only motifs occurring in 5% or more of foreground regions in HOMER results were considered. The R package ggplot [59] was used to produce bubble plots for network visualization, where negative log p-value determine bubble size.

### Bulk RNA-seq

RNA was isolated from whole EBs using Trizol (Invitrogen). RNA was isolated from two independent MEIS KO clones, both of which exhibited the same phenotypic response (lack of NKX2-5/GFP expression). For each MEIS KO clone, RNA was extracted at d5 and d7 in the course of two independent differentiation (total of four differentiations, each including a wild-type control). The quality and integrity of the RNA samples was assessed using a 4200 TapeStation (Agilent Technologies) and libraries generated using the Illumina® Stranded mRNA Prep Ligation kit (Illumina, Inc.) according to the manufacturer’s protocol. The loaded flow-cell was paired-end sequenced (76 + 76 cycles, plus indices) on an Illumina HiSeq4000 instrument. RNA-seq experiments were analysed using Trimmomatic for trimming [60], STAR [61] for aligning to the human genome (hg38). DESeq2 [62] was used to normalize the gene expression count values, calculate fold changes, and to detect the differential expression genes across WT and KO using adjusted p-value < 0.05 and log FC 1 as the cutoff values. Enrichr [63] was used to determine functional enrichment of the differential expressed genes. For the analysis of genes encoding for DE TFs, human TFs were downloaded from [64] and only TFs associated with a MEIS peak using GREAT standard association rule settings [17] were used.

### ChIP-seq

ChIPmentation assays were performed as described [65], starting from 3-5x10^6^ cell for TFs and 0.5-1x10^6^ for H3K27ac and using the following antibodies: MEIS1( Abcam ab19867), GATA-6 XP Rabbit mAb (D61E4, Cell Signaling Technology), H3K27ac (Abcam ab4729), rabbit polyclonal KMT2D (Atlas Antibodies) and monoclonal ANTI-FLAG M2 antibody (Sigma-Aldrich). ChIP-seq experiments were analysed using Trimmomatic for trimming [60], Bowtie2 [66] for aligning to the human genome (hg38), SAMtools [67] to remove the aligned reads with a mapping quality Q30 and MACS2 for peak calling [68] with default narrow peak calling setting for TFs. Only high confidence binding regions, fold enrichment (FE) > 10, were considered for target assignments and GO analysis. TF ChIP-seq were also analysed with the Bioconductor package Motif2Site [69], as specified in the relative figure legend. Motif2Site detects TF binding sites by performing peak calling around user-provided motif sets and re-centers binding sites across replicate ChIP-seq experiments to improve the prediction accuracy. ‘*findMotifGenome’* module of the HOMER package [22] was used to detect *de novo* motif in 200nt summit regions. TFs motifs along with LOD values are listed in Table S4. For H3K27ac and KMT2D ChIP-seq, short reads were counted in open chromatin regions from ATAC-seq. Regions of size 3000 and 1000 nucleotides for H3K27ac and KMT2D, respectively, were centred on open chromatin peaks. Rsubread [70] was used to generate count values and EdgeR [71] was used to normalize count values and calculate fold changes of all ChIP-seq experiments across WT and MEIS KO. GREAT standard association rule settings [17] was used to associate ChIP-seq peaks with genes and uncover events controlled by TF binding. ggplot2 package [59] was used to generate all the used plots, GenomicRanges package [72] was used to perform various operation on genomic intervals, and eulerr was used to generate Venn diagrams [73].

### ATAC-seq sequencing data and downstream analyses

Cell pellets lysis, tagmentation and DNA purification were performed on 50-100x10^3^ cells/sample using ATAC-Seq Kit (Active Motif) following manufacturer’s instructions. ATAC-seq experiments were analysed using Trimmomatic for trimming [60], Bowtie2 [66] for aligning to the human genome (hg38), SAMtools [67] to remove the aligned reads with a mapping quality Q30 and MACS2 for peak calling [68] with parameter setting for ATAC-seq peak calling. DiffBind [74] was used to re-center peaks across the called peaks and generated the count data in those peaks. EdgeR [71] was used to normalize count values, calculate fold changes, and detect differential open chromatin regions using FDR 0.1 as cutoff value. ‘*findMotifGenome’* module of the HOMER package was used to detect *de novo* motif in differential accessible regions [22]. GREAT standard association rule settings [17] was used to annotate differential accessible regions across WT and MEIS KO.

### Single cell RNA-seq and downstream analyses

Differentiations were started on sequential days so the different timepoints could be collected and processed on the same day. Sample 1 was collected on day 3 and day 3.75 (50:50 mix); Samples 2, 3 and 4 were collected on days 4.75, 5.75 and 12 respectively. EBs were dissociated to single cells using TrypLE Select. Gene expression libraries were prepared from the single cells using the Chromium Controller and Single Cell 3ʹ Reagent Kits v3 (10x Genomics) according to the manufacturer’s protocol (CG000183 Rev A). Sequencing libraries comprised standard Illumina paired-end constructs. Paired-end sequencing (26:98) was performed on the Illumina NextSeq500 platform using NextSeq 500/550 High Output v2.5 (150 Cycles) reagents. The .bcl sequence data were processed for QC purposes using bcl2fastq software (v. 2.20.0.422) and the resulting .fastq files assessed using FastQC (v. 0.11.3), FastqScreen (v. 0.9.2) and FastqStrand (v. 0.0.7) prior to pre-processing with the CellRanger pipeline. Raw sequencing data were processed using the 10x Genomics Cell Ranger pipeline (v3.1.0). Base call (BCL) files generated by the sequencer were converted to FASTQ files using “cellranger mkfastq”. The FASTQ files were mapped against the pre-built human reference package from 10X Genomics (GRCh38-3.0.0). After removing outlier cells (total read counts >40,000) 4,860 cells (3,509 cells from day 3 and 3.75; 5013 cells from day 4.75; 3,984 cells from day 5.75; and 2,124 cells from day 12) remained for downstream analysis. We filtered out genes with average UMI counts per cell below 0.01. To account for differences in sequencing depth per cell, we normalized the raw counts using a deconvolution-based method and then log-transformed with a pseudo-count of 1 added to all counts. The QC filtered cells from the different timepoints were aggregated for downstream analysis. Highly Variable Genes (HVGs) were identified using the modelGeneVar function from scran [75], using biological components >0.5 and FDR value <0.1. Cell cycle phase classification was performed using the cyclone function from scran and after evaluating their impact HVGs strongly associated with cell cycle were subtracted. These HVGs were then used to reduce the dimensions of the dataset using PCA and to 2D using t-SNE, where the first 14 components of the PCA were given as input. Cells were grouped into clusters using the dynamicTreeCut package [76]. Clusters were assigned to cell types using known marker genes.

## Data availability

All data described have been deposited in ArrayExpress with the following accession numbers:

E-MTAB-14241: Replicate HOXB1 ChIP-seq

E-MTAB-14243: Replicate ChIP-seq for MEIS1; GATA6 (wild-type and KO); H3K27Ac (wild-type and KO); KMT2D (wild-type and KO).

E-MTAB-14242: Replicate ATAC-seq (wild-type and KO; day5 and day12).

E-MTAB-14250: RNA-seq (wild-type and MEIS KO cardiac differentiation).

E-MTAB-14248: RNA-seq (timecourse of hESC cardiac differentiation).

E-MTAB-14249: ScRNA-seq (timecourse of hESC cardiac differentiation).

---

**Figure S1.**
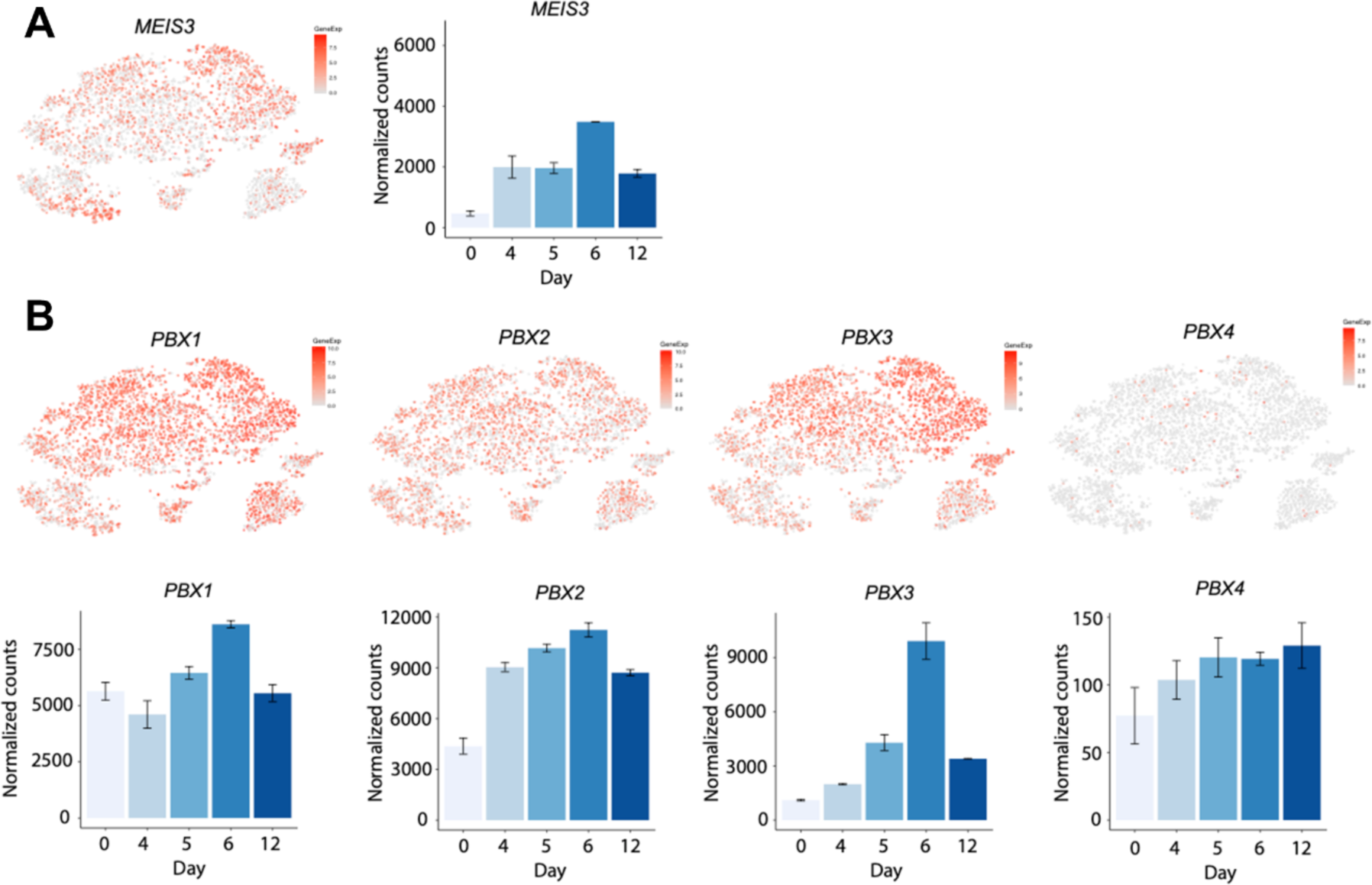
AB. t-SNE plots showing gene expression level in the scRNA-seq cardiac differentiation time-course, and corresponding bulk RNA-seq normalized counts by differentiation day, for *MEIS3* (A) and PBX family members (B). The scRNA-seq cell populations are annotated in Figure 1D.

**Figure S2.**
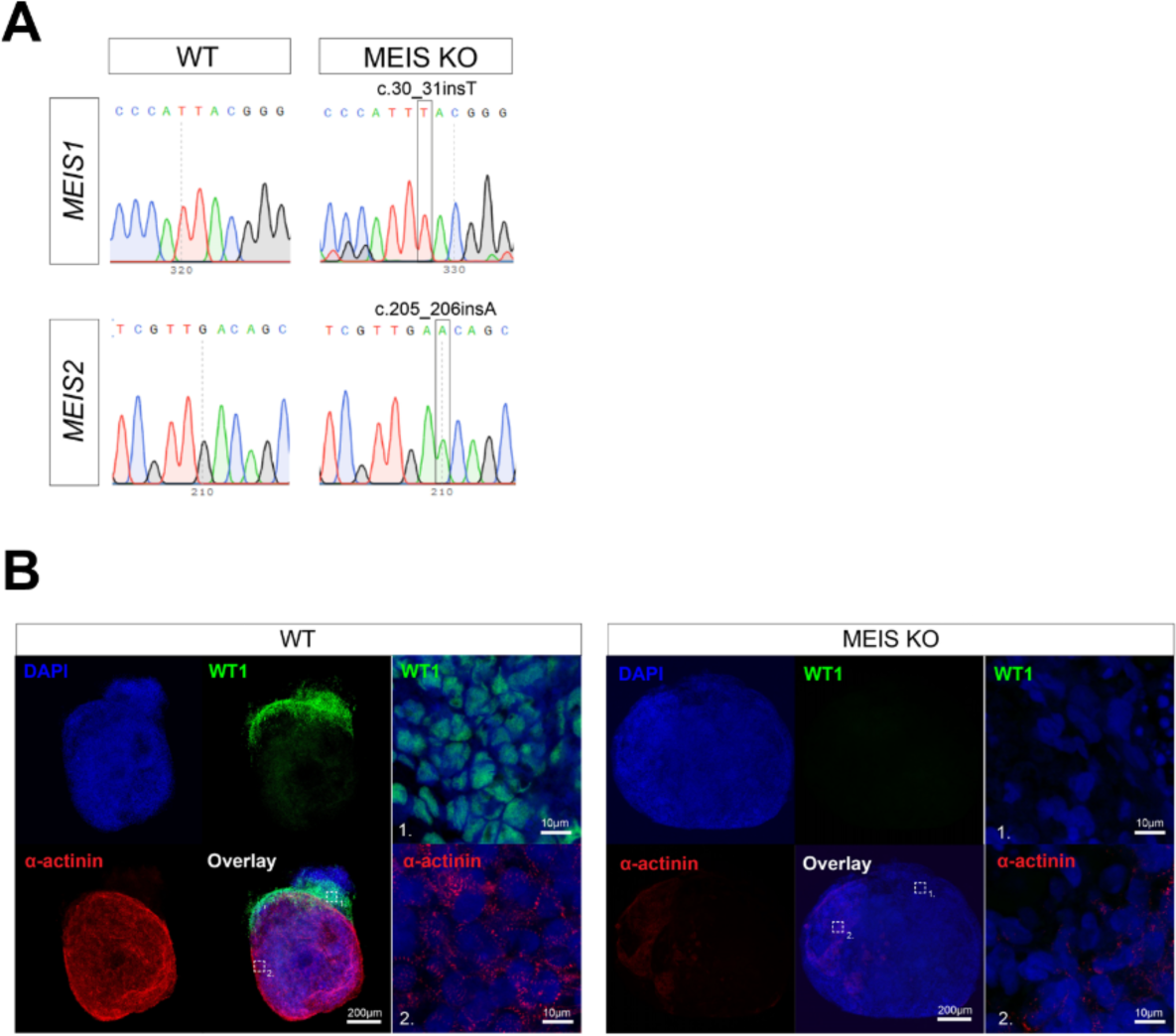
A. Sanger sequencing chromatograms showing the presence of homozygous insertion frameshift-causing mutations in *MEIS1* (c.30_31insT) and *MEIS2* (c.205_206insA) of the MEIS KO clone. The reference wild-type sequencing is also shown. B. Confocal microscopy image of WT1 (epicardial cells, green), α-actinin (cardiomyocytes, red) and nuclei (DAPI, blue) in day 12 EBs generated from wild-type and MEISKO hESCs. The high magnification inserts show a sarcomere staining pattern with α-actinin and nuclear WT1, both of which are absent in the KO line.

**Figure S3.**
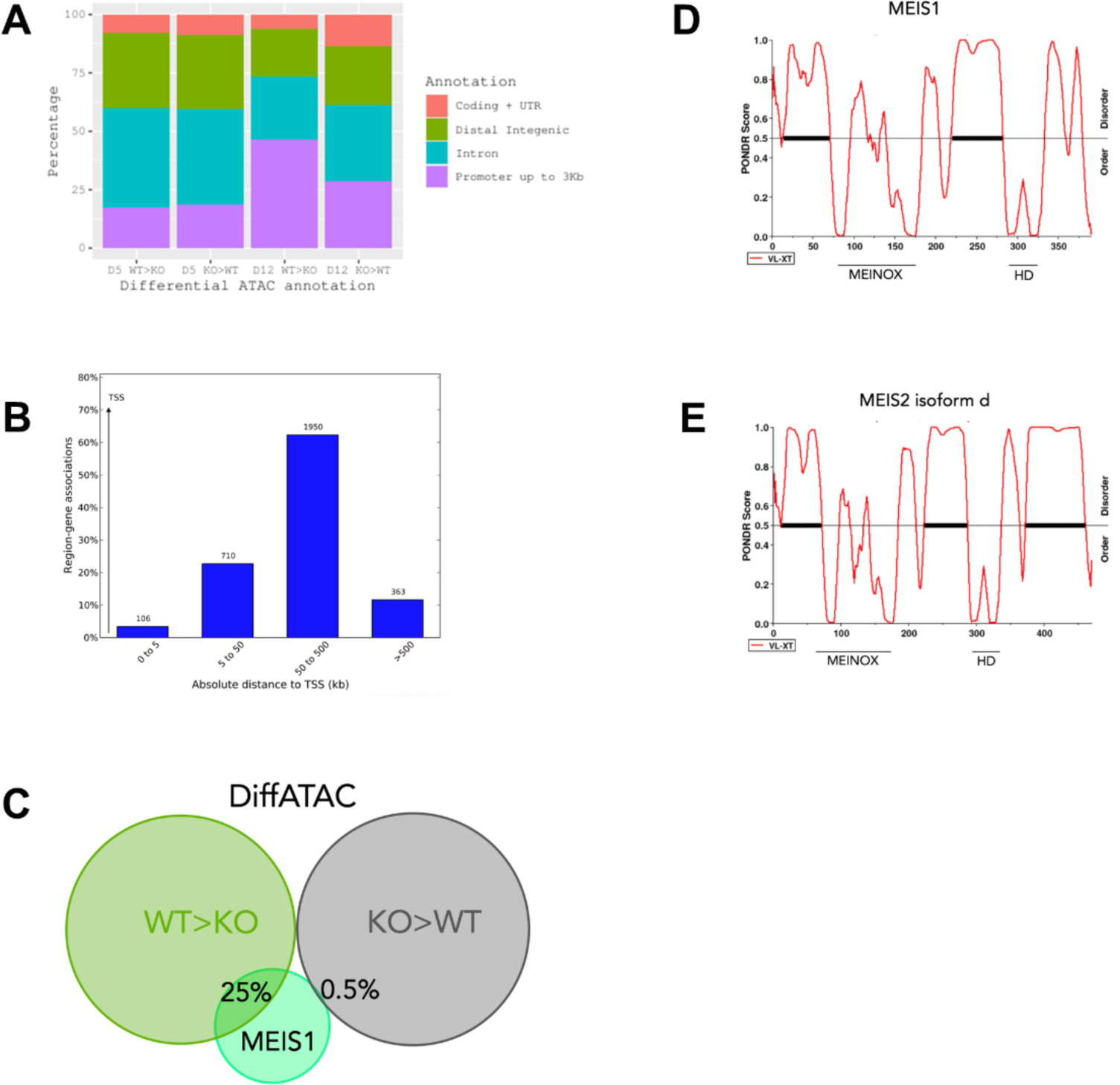
A. Genomic distribution of differential chromatin accessibility in wild-type and MEIS1-2 KO at d5 and d12. At d5, differential accessibility between wild-type and mutant is largely observed at distal regions and introns. At d12, promoters make half of the differentially accessible regions in wild-type while in the absence of MEIS1-2, distal regions and introns remain more highly accessible. B. Genomic distribution of high-confidence MEIS1 peaks. C. Intersection of high-confidence MEIS1 peaks with differential ATAC-seq peaks. MEIS1 binding overlaps almost exclusively with chromatin that is more accessible in wild-type conditions (25% of top MEIS1 peaks; n=446), indicating a direct association between MEIS1 binding and accessible chromatin. In contrast, in the mutant, chromatin accessibility is independent of MEIS, with only 0.5% of top MEIS peaks (n=8) intersecting ATAC peaks higher in KO. This suggests that MEIS does not directly establish repressive chromatin environments; rather, sites that become more accessible in the absence of MEIS are likely opened by upregulated TFs. D, E. Output of PONDR (http://www.pondr.com) [77] for MEIS1 and MEIS2 (isoform d). MEIS1-2 contain a majority of IDRs and two well-conserved structured domains, the MEINOX domain and the homeodomain [9]. The disordered C terminal domain contains MEIS1 activation domain [78,79].

**Figure S4.**
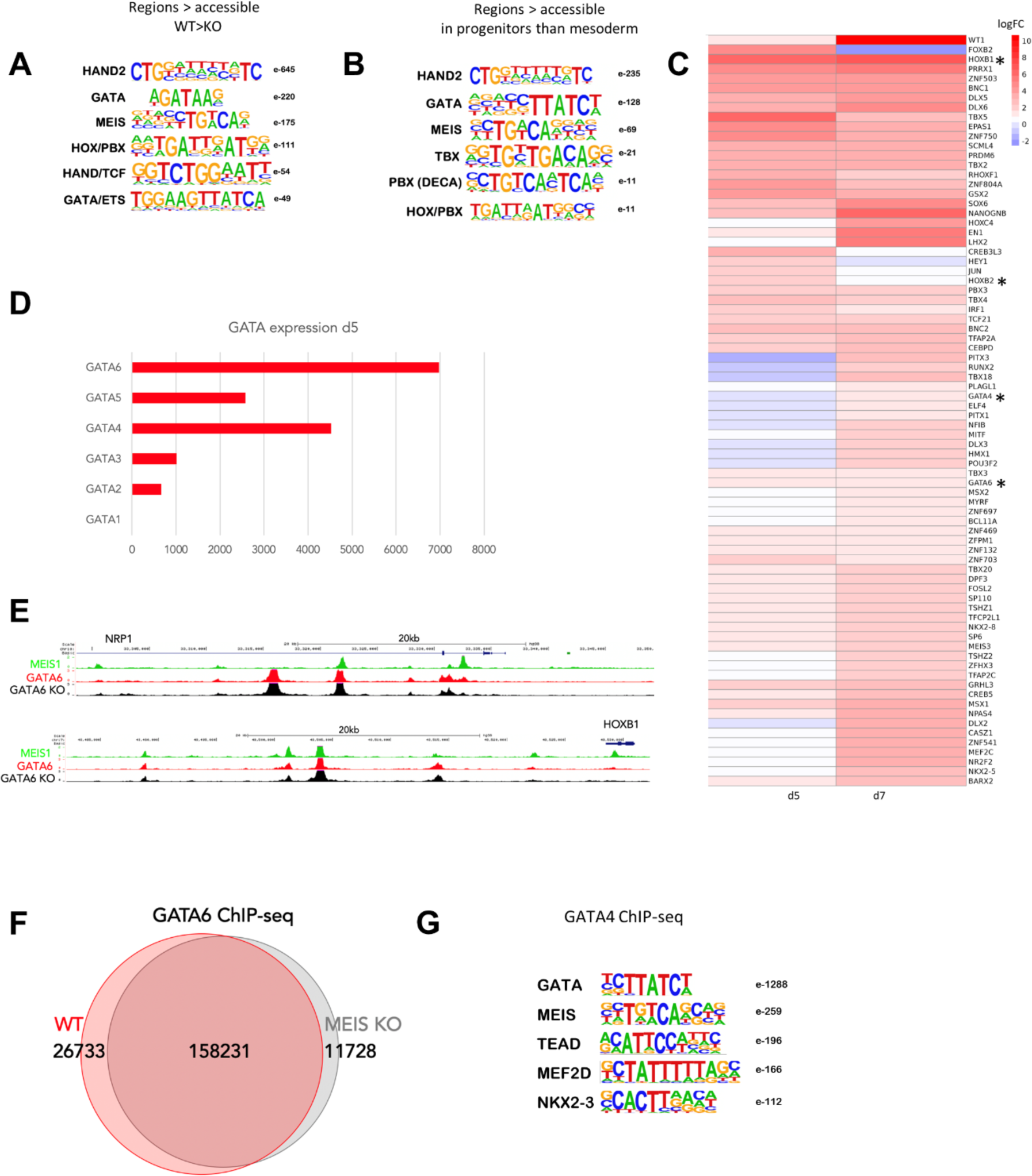
A. Top motifs, ranked by significance, identified in differential ATAC WT>KO (regions with lower accessibility in MEIS KO) using de novo motif discovery (Homer). B. Top motifs, ranked by significance, identified in regions that increase accessibility in cardiac progenitors relative to mesoderm; data sourced from [15] using de novo motif discovery (Homer). C. Differentially expressed TFs in MEIS KO. Log FC WT versus MEIS KO at d5 and d7. D. Normalised RNA-seq counts for GATA family members in d5 cardiac progenitors. E. UCSC tracks of MEIS1 (green) and GATA6 in wild-type and MEIS KO cells (red and black, respectively) at the *NRP1* and *HOXB1* loci. F. Intersection of GATA6 binding in d5 WT and MEIS KO progenitors. The Venn diagram was generated using all statistically significant GATA6 peaks identified using Motif2site [69]. G. Top motifs, ranked by significance, enriched in GATA4 ChIP-seq in d6 iPSCs-derived cardiac progenitors; data sourced from [26] using de novo motif discovery (Homer).

**Figure S5.**
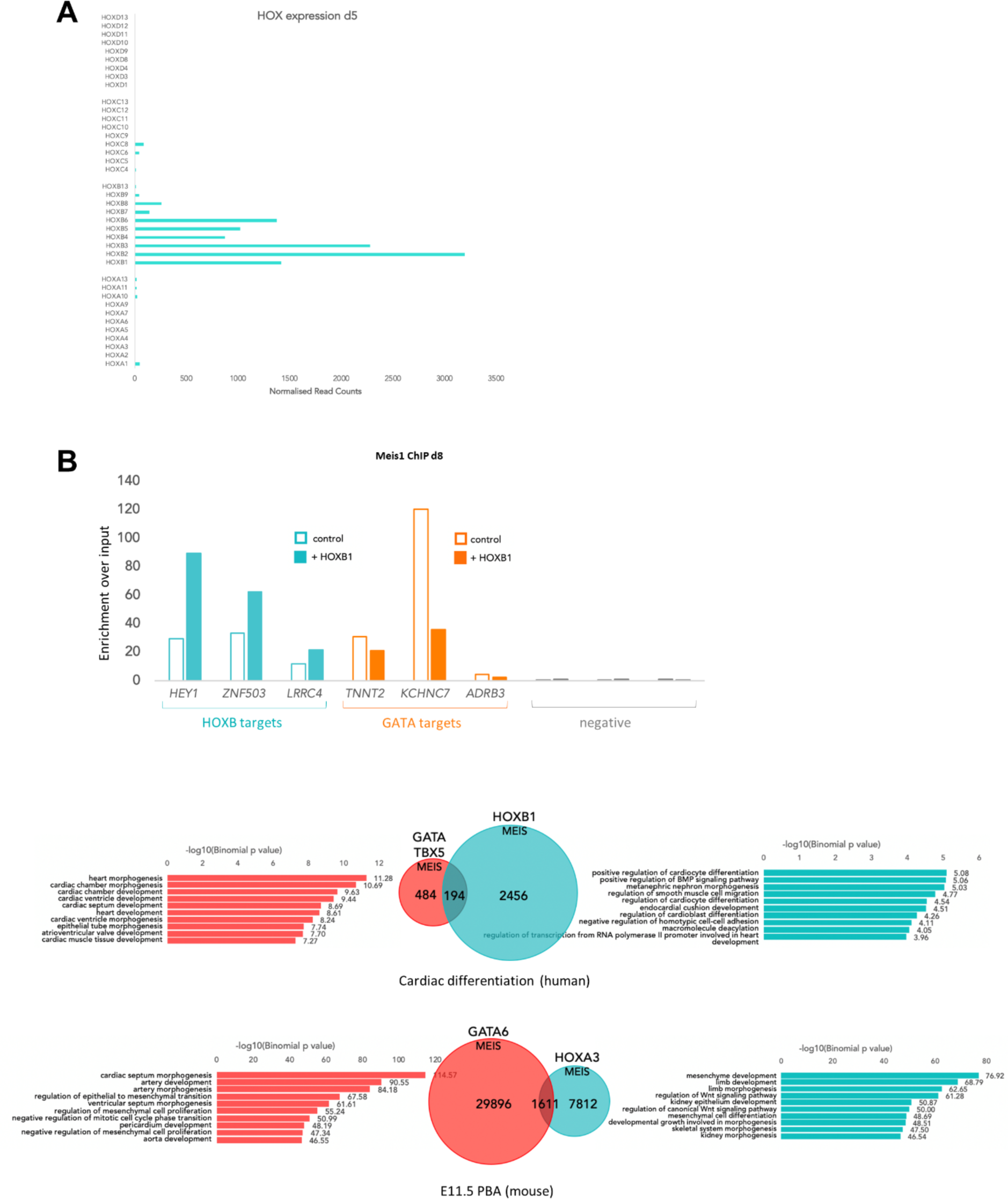
A. Normalised RNA-seq counts for HOX family members in d5 cardiac progenitors. B. One of the experiments shown in Fig 5G, where MEIS enrichment over input is measured at HOXB1 and GATA target enhancers in d8 control and HOXB1-overexpressing cells. Negative control loci are regions with no detectable MEIS1 binding signal, located in the vicinity of *HOXC11*, *AMO3* and *SMAD6*. C. Overlap between MEIS-HOX and MEIS-GATA co-occupied regions. To assess the full extent of the overlap, MEIS-HOX and MEIS-GATA co-bound regions were detected on a genome-wide scale using Motif2Site [69]. The set of GATA regions included those co-occupied by GATA6 (this paper) and GATA4-TBX5 regions [26] in cardiac progenitors. GATA-MEIS only regions are specifically associated with cardiac differentiation GO terms, while HOX-MEIS only regions are associated with regulatory GO terms, which include regulation of cardiac differentiation as well as regulation of other processes. D. Overlap of GATA6, MEIS and HOXA3 binding in the cardiac neural crest-populated posterior branchial arches (PBA) [31]. Similar to human cardiac differentiation, GATA6-MEIS binding is highly cardiac-specific while MEIS-HOX is linked with diverse developmental processes.

## References

1. Spitz, F.; Furlong, E.E.M. Transcription factors: from enhancer binding to developmental control. Nature Reviews Genetics 2012, 13, 613–626, doi:10.1038/nrg3207.

2. Wang, H.; Yang, Y.; Liu, J.; Qian, L. Direct cell reprogramming: approaches, mechanisms and progress. Nat Rev Mol Cell Biol 2021, 22, 410–424, doi:10.1038/s41580-021-00335-z.

3. Reiter, F.; Wienerroither, S.; Stark, A. Combinatorial function of transcription factors and cofactors. Curr Opin Genet Dev 2017, 43, 73–81, doi:10.1016/j.gde.2016.12.007.

4. Mirny, L.A. Nucleosome-mediated cooperativity between transcription factors. Proc Natl Acad Sci U S A 2010, 107, 22534–22539, doi:10.1073/pnas.0913805107.

5. Moyle-Heyrman, G.; Tims, H.S.; Widom, J. Structural constraints in collaborative competition of transcription factors against the nucleosome. J Mol Biol 2011, 412, 634–646, doi:10.1016/j.jmb.2011.07.032.

6. Selleri, L.; Zappavigna, V.; Ferretti, E. ’Building a perfect body’: control of vertebrate organogenesis by PBX-dependent regulatory networks. Genes Dev 2019, 33, 258–275, doi:10.1101/gad.318774.118.

7. Bürglin, T.R.; Affolter, M. Homeodomain proteins: an update. Chromosoma 2016, 125, 497–521, doi:10.1007/s00412-015-0543-8.

8. Bobola, N.; Sagerström, C.G. TALE transcription factors: Cofactors no more. Seminars in Cell & Developmental Biology 2024, 152-153, 76-84, 10.1016/j.semcdb.2022.11.015.

9. Schulte, D.; Geerts, D. MEIS transcription factors in development and disease. Development 2019, 146, doi:10.1242/dev.174706.

10. Bridoux, L.; Zarrineh, P.; Mallen, J.; Phuycharoen, M.; Latorre, V.; Ladam, F.; Losa, M.; Baker, S.M.; Sagerstrom, C.; Mace, K.A.;, et al. HOX paralogs selectively convert binding of ubiquitous transcription factors into tissue-specific patterns of enhancer activation. PLOS Genetics 2020, 16, e1009162, doi:10.1371/journal.pgen.1009162.

11. Phuycharoen, M.; Zarrineh, P.; Bridoux, L.; Amin, S.; Losa, M.; Chen, K.; Bobola, N.; Rattray, M. Uncovering tissue-specific binding features from differential deep learning. Nucleic Acids Research 2020, 48, e27–e27, doi:10.1093/nar/gkaa009.

12. Garcia-Mora, A.; Mallen, J.; Zarrineh, P.; Hanley, N.; Gerrard, D.; Bobola, N. A pipeline to identify TF combinatorial binding uncovers TEAD1 as an antagonist of tissue-specific transcription factors in human organogenesis. bioRxiv 2023, 2023.2010.2005.561094, doi:10.1101/2023.10.05.561094.

13. Birket, M.J.; Ribeiro, M.C.; Verkerk, A.O.; Ward, D.; Leitoguinho, A.R.; den Hartogh, S.C.; Orlova, V.V.; Devalla, H.D.; Schwach, V.; Bellin, M.;, et al. Expansion and patterning of cardiovascular progenitors derived from human pluripotent stem cells. Nat Biotechnol 2015, 33, 970–979, doi:10.1038/nbt.3271.

14. Elliott, D.A.; Braam, S.R.; Koutsis, K.; Ng, E.S.; Jenny, R.; Lagerqvist, E.L.; Biben, C.; Hatzistavrou, T.; Hirst, C.E.; Yu, Q.C.;, et al. NKX2-5eGFP/w hESCs for isolation of human cardiac progenitors and cardiomyocytes. Nature Methods 2011, 8, 1037–1040, doi:10.1038/nmeth.1740.

15. Bertero, A.; Fields, P.A.; Ramani, V.; Bonora, G.; Yardimci, G.G.; Reinecke, H.; Pabon, L.; Noble, W.S.; Shendure, J.; Murry, C.E. Dynamics of genome reorganization during human cardiogenesis reveal an RBM20-dependent splicing factory. Nature Communications 2019, 10, 1538, doi:10.1038/s41467-019-09483-5.

16. Wu, X.; Wu, X.; Xie, W. Activation, decommissioning, and dememorization: enhancers in a life cycle. Trends in Biochemical Sciences 2023, 48, 673–688, 10.1016/j.tibs.2023.04.005.

17. McLean, C.Y.; Bristor, D.; Hiller, M.; Clarke, S.L.; Schaar, B.T.; Lowe, C.B.; Wenger, A.M.; Bejerano, G. GREAT improves functional interpretation of cis-regulatory regions. Nat Biotechnol 2010, 28, 495–501, doi:10.1038/nbt.1630.

18. Klemm, S.L.; Shipony, Z.; Greenleaf, W.J. Chromatin accessibility and the regulatory epigenome. Nature Reviews Genetics 2019, 20, 207–220, doi:10.1038/s41576-018-0089-8.

19. Creyghton, M.P.; Cheng, A.W.; Welstead, G.G.; Kooistra, T.; Carey, B.W.; Steine, E.J.; Hanna, J.; Lodato, M.A.; Frampton, G.M.; Sharp, P.A.;, et al. Histone H3K27ac separates active from poised enhancers and predicts developmental state. Proc Natl Acad Sci U S A 2010, 107, 21931–21936, doi:10.1073/pnas.1016071107.

20. Boija, A.; Klein, I.A.; Sabari, B.R.; Dall’Agnese, A.; Coffey, E.L.; Zamudio, A.V.; Li, C.H.; Shrinivas, K.; Manteiga, J.C.; Hannett, N.M.;, et al. Transcription Factors Activate Genes through the Phase-Separation Capacity of Their Activation Domains. Cell 2018, 175, 1842–1855.e1816, doi:10.1016/j.cell.2018.10.042.

21. Sabari, B.R.; Dall’Agnese, A.; Boija, A.; Klein, I.A.; Coffey, E.L.; Shrinivas, K.; Abraham, B.J.; Hannett, N.M.; Zamudio, A.V.; Manteiga, J.C.;, et al. Coactivator condensation at super-enhancers links phase separation and gene control. Science 2018, 361, doi:10.1126/science.aar3958.

22. Heinz, S.; Benner, C.; Spann, N.; Bertolino, E.; Lin, Y.C.; Laslo, P.; Cheng, J.X.; Murre, C.; Singh, H.; Glass, C.K. Simple combinations of lineage-determining transcription factors prime cis-regulatory elements required for macrophage and B cell identities. Mol Cell 2010, 38, 576–589, doi:10.1016/j.molcel.2010.05.004.

23. Tremblay, M.; Sanchez-Ferras, O.; Bouchard, M. GATA transcription factors in development and disease. Development 2018, 145, doi:10.1242/dev.164384.

24. Sam, J.; Mercer, E.J.; Torregroza, I.; Banks, K.M.; Evans, T. Specificity, redundancy and dosage thresholds among gata4/5/6 genes during zebrafish cardiogenesis. Biol Open 2020, 9, doi:10.1242/bio.053611.

25. Xin, M.; Davis, C.A.; Molkentin, J.D.; Lien, C.-L.; Duncan, S.A.; Richardson, J.A.; Olson, E.N. A threshold of *GATA4* and *GATA6* expression is required for cardiovascular development. Proceedings of the National Academy of Sciences 2006, 103, 11189-11194, doi:doi:10.1073/pnas.0604604103.

26. Gonzalez-Teran, B.; Pittman, M.; Felix, F.; Thomas, R.; Richmond-Buccola, D.; Hüttenhain, R.; Choudhary, K.; Moroni, E.; Costa, M.W.; Huang, Y.;, et al. Transcription factor protein interactomes reveal genetic determinants in heart disease. Cell 2022, 185, 794–814.e730, doi:10.1016/j.cell.2022.01.021.

27. Rezsohazy, R.; Saurin, A.J.; Maurel-Zaffran, C.; Graba, Y. Cellular and molecular insights into Hox protein action. Development 2015, 142, 1212–1227, doi:10.1242/dev.109785.

28. Stefanovic, S.; Laforest, B.; Desvignes, J.-P.; Lescroart, F.; Argiro, L.; Maurel-Zaffran, C.; Salgado, D.; Plaindoux, E.; De Bono, C.; Pazur, K.;, et al. Hox-dependent coordination of mouse cardiac progenitor cell patterning and differentiation. eLife 2020, 9, e55124, doi:10.7554/eLife.55124.

29. Zhu, F.; Gamboa, M.; Farruggio, A.P.; Hippenmeyer, S.; Tasic, B.; Schüle, B.; Chen-Tsai, Y.; Calos, M.P. DICE, an efficient system for iterative genomic editing in human pluripotent stem cells. Nucleic Acids Res 2014, 42, e34, doi:10.1093/nar/gkt1290.

30. Ieda, M.; Fu, J.D.; Delgado-Olguin, P.; Vedantham, V.; Hayashi, Y.; Bruneau, B.G.; Srivastava, D. Direct reprogramming of fibroblasts into functional cardiomyocytes by defined factors. Cell 2010, 142, 375–386, doi:10.1016/j.cell.2010.07.002.

31. Losa, M.; Latorre, V.; Andrabi, M.; Ladam, F.; Sagerström, C.; Novoa, A.; Zarrineh, P.; Bridoux, L.; Hanley, N.A.; Mallo, M.; Bobola, N. A tissue-specific, Gata6-driven transcriptional program instructs remodeling of the mature arterial tree. eLife 2017, 6, e31362, doi:10.7554/eLife.31362.

32. Gangfuß, A.; Yigit, G.; Altmüller, J.; Nürnberg, P.; Czeschik, J.C.; Wollnik, B.; Bögershausen, N.; Burfeind, P.; Wieczorek, D.; Kaiser, F.;, et al. Intellectual disability associated with craniofacial dysmorphism, cleft palate, and congenital heart defect due to a de novo MEIS2 mutation: A clinical longitudinal study. Am J Med Genet A 2021, 185, 1216–1221, doi:10.1002/ajmg.a.62070.

33. Ng, S.B.; Bigham, A.W.; Buckingham, K.J.; Hannibal, M.C.; McMillin, M.J.; Gildersleeve, H.I.; Beck, A.E.; Tabor, H.K.; Cooper, G.M.; Mefford, H.C.;, et al. Exome sequencing identifies MLL2 mutations as a cause of Kabuki syndrome. Nature Genetics 2010, 42, 790–793, doi:10.1038/ng.646.

34. Hu, D.; Gao, X.; Morgan, M.A.; Herz, H.M.; Smith, E.R.; Shilatifard, A. The MLL3/MLL4 branches of the COMPASS family function as major histone H3K4 monomethylases at enhancers. Mol Cell Biol 2013, 33, 4745–4754, doi:10.1128/mcb.01181-13.

35. Froimchuk, E.; Jang, Y.; Ge, K. Histone H3 lysine 4 methyltransferase KMT2D. Gene 2017, 627, 337–342, doi:10.1016/j.gene.2017.06.056.

36. Ang, S.Y.; Uebersohn, A.; Spencer, C.I.; Huang, Y.; Lee, J.E.; Ge, K.; Bruneau, B.G. KMT2D regulates specific programs in heart development via histone H3 lysine 4 di-methylation. Development 2016, 143, 810–821, doi:10.1242/dev.132688.

37. Aubrey, B.J.; Cutler, J.A.; Bourgeois, W.; Donovan, K.A.; Gu, S.; Hatton, C.; Perlee, S.; Perner, F.; Rahnamoun, H.; Theall, A.C.P.;, et al. IKAROS and MENIN coordinate therapeutically actionable leukemogenic gene expression in MLL-r acute myeloid leukemia. Nature Cancer 2022, 3, 595–613, doi:10.1038/s43018-022-00366-1.

38. Lee, J.-E.; Wang, C.; Xu, S.; Cho, Y.-W.; Wang, L.; Feng, X.; Baldridge, A.; Sartorelli, V.; Zhuang, L.; Peng, W.; Ge, K. H3K4 mono- and di-methyltransferase MLL4 is required for enhancer activation during cell differentiation. eLife 2013, 2, e01503, doi:10.7554/eLife.01503.

39. Lai, B.; Lee, J.E.; Jang, Y.; Wang, L.; Peng, W.; Ge, K. MLL3/MLL4 are required for CBP/p300 binding on enhancers and super-enhancer formation in brown adipogenesis. Nucleic Acids Res 2017, 45, 6388–6403, doi:10.1093/nar/gkx234.

40. Mukherjee, K.; Bürglin, T.R. Comprehensive Analysis of Animal TALE Homeobox Genes: New Conserved Motifs and Cases of Accelerated Evolution. Journal of Molecular Evolution 2007, 65, 137–153, 10.1007/s00239-006-0023-0.

41. Joo, S.; Wang, M.H.; Lui, G.; Lee, J.; Barnas, A.; Kim, E.; Sudek, S.; Worden, A.Z.; Lee, J.H. Common ancestry of heterodimerizing TALE homeobox transcription factors across Metazoa and Archaeplastida. BMC Biol 2018, 16, 136, doi:10.1186/s12915-018-0605-5.

42. Hoffman, J.I.; Kaplan, S. The incidence of congenital heart disease. J Am Coll Cardiol 2002, 39, 1890–1900, doi:10.1016/s0735-1097(02)01886-7.

43. Cui, M.; Wang, Z.; Bassel-Duby, R.; Olson, E.N. Genetic and epigenetic regulation of cardiomyocytes in development, regeneration and disease. Development 2018, 145, doi:10.1242/dev.171983.

44. Paige, Sharon L.; Thomas, S.; Stoick-Cooper, Cristi L.; Wang, H.; Maves, L.; Sandstrom, R.; Pabon, L.; Reinecke, H.; Pratt, G.; Keller, G.;, et al. A Temporal Chromatin Signature in Human Embryonic Stem Cells Identifies Regulators of Cardiac Development. Cell 2012, 151, 221–232, 10.1016/j.cell.2012.08.027.

45. Wamstad, J.A.; Alexander, J.M.; Truty, R.M.; Shrikumar, A.; Li, F.; Eilertson, K.E.; Ding, H.; Wylie, J.N.; Pico, A.R.; Capra, J.A.;, et al. Dynamic and coordinated epigenetic regulation of developmental transitions in the cardiac lineage. Cell 2012, 151, 206–220, doi:10.1016/j.cell.2012.07.035.

46. Dupays, L.; Shang, C.; Wilson, R.; Kotecha, S.; Wood, S.; Towers, N.; Mohun, T. Sequential Binding of MEIS1 and NKX2-5 on the Popdc2 Gene: A Mechanism for Spatiotemporal Regulation of Enhancers during Cardiogenesis. Cell Reports 2015, 13, 183–195, 10.1016/j.celrep.2015.08.065.

47. Bouilloux, F.; Thireau, J.; Ventéo, S.; Farah, C.; Karam, S.; Dauvilliers, Y.; Valmier, J.; Copeland, N.G.; Jenkins, N.A.; Richard, S.; Marmigère, F. Loss of the transcription factor Meis1 prevents sympathetic neurons target-field innervation and increases susceptibility to sudden cardiac death. eLife 2016, 5, e11627, doi:10.7554/eLife.11627.

48. Quaranta, R.; Fell, J.; Rühle, F.; Rao, J.; Piccini, I.; Araúzo-Bravo, M.J.; Verkerk, A.O.; Stoll, M.; Greber, B. Revised roles of ISL1 in a hES cell-based model of human heart chamber specification. eLife 2018, 7, e31706, doi:10.7554/eLife.31706.

49. Delgado, I.; Giovinazzo, G.; Temiño, S.; Gauthier, Y.; Balsalobre, A.; Drouin, J.; Torres, M. Control of mouse limb initiation and antero-posterior patterning by Meis transcription factors. Nature Communications 2021, 12, 3086, doi:10.1038/s41467-021-23373-9.

50. Luna-Zurita, L.; Stirnimann, C.U.; Glatt, S.; Kaynak, B.L.; Thomas, S.; Baudin, F.; Samee, M.A.; He, D.; Small, E.M.; Mileikovsky, M.;, et al. Complex Interdependence Regulates Heterotypic Transcription Factor Distribution and Coordinates Cardiogenesis. Cell 2016, 164, 999–1014, doi:10.1016/j.cell.2016.01.004.

51. Garg, V.; Kathiriya, I.S.; Barnes, R.; Schluterman, M.K.; King, I.N.; Butler, C.A.; Rothrock, C.R.; Eapen, R.S.; Hirayama-Yamada, K.; Joo, K.;, et al. GATA4 mutations cause human congenital heart defects and reveal an interaction with TBX5. Nature 2003, 424, 443–447, doi:10.1038/nature01827.

52. Amin, S.; Donaldson, Ian J.; Zannino, D.A.; Hensman, J.; Rattray, M.; Losa, M.; Spitz, F.; Ladam, F.; Sagerström, C.; Bobola, N. Hoxa2 Selectively Enhances Meis Binding to Change a Branchial Arch Ground State. Developmental Cell 2015, 32, 265–277, 10.1016/j.devcel.2014.12.024.

53. Dvoretskova, E.; Ho, M.C.; Kittke, V.; Neuhaus, F.; Vitali, I.; Lam, D.D.; Delgado, I.; Feng, C.; Torres, M.; Winkelmann, J.; Mayer, C. Spatial enhancer activation influences inhibitory neuron identity during mouse embryonic development. Nature Neuroscience 2024, 27, 862–872, doi:10.1038/s41593-024-01611-9.

54. Dupacova, N.; Antosova, B.; Paces, J.; Kozmik, Z. Meis homeobox genes control progenitor competence in the retina. Proc Natl Acad Sci U S A 2021, 118, doi:10.1073/pnas.2013136118.

55. Ji, Z.; Donaldson, I.J.; Liu, J.; Hayes, A.; Zeef, L.A.; Sharrocks, A.D. The forkhead transcription factor FOXK2 promotes AP-1-mediated transcriptional regulation. Mol Cell Biol 2012, 32, 385–398, doi:10.1128/mcb.05504-11.

56. Gerrard, D.T.; Berry, A.A.; Jennings, R.E.; Piper Hanley, K.; Bobola, N.; Hanley, N.A. An integrative transcriptomic atlas of organogenesis in human embryos. eLife 2016, 5, e15657, doi:10.7554/eLife.15657.

57. Gerrard, D.T.; Berry, A.A.; Jennings, R.E.; Birket, M.J.; Zarrineh, P.; Garstang, M.G.; Withey, S.L.; Short, P.; Jiménez-Gancedo, S.; Firbas, P.N.;, et al. Dynamic changes in the epigenomic landscape regulate human organogenesis and link to developmental disorders. Nat Commun 2020, 11, 3920, doi:10.1038/s41467-020-17305-2.

58. Quinlan, A.R.; Hall, I.M. BEDTools: a flexible suite of utilities for comparing genomic features. Bioinformatics 2010, 26, 841–842, doi:10.1093/bioinformatics/btq033.

59. Wickham, H. ggplot2: Elegant Graphics for Data Analysis; Springer-Verlag New York, 2016.

60. Bolger, A.M.; Lohse, M.; Usadel, B. Trimmomatic: a flexible trimmer for Illumina sequence data. Bioinformatics 2014, 30, 2114–2120, doi:10.1093/bioinformatics/btu170.

61. Dobin, A.; Davis, C.A.; Schlesinger, F.; Drenkow, J.; Zaleski, C.; Jha, S.; Batut, P.; Chaisson, M.; Gingeras, T.R. STAR: ultrafast universal RNA-seq aligner. Bioinformatics 2013, 29, 15–21, doi:10.1093/bioinformatics/bts635.

62. Love, M.I.; Huber, W.; Anders, S. Moderated estimation of fold change and dispersion for RNA-seq data with DESeq2. Genome Biology 2014, 15, 550, doi:10.1186/s13059-014-0550-8.

63. Kuleshov, M.V.; Jones, M.R.; Rouillard, A.D.; Fernandez, N.F.; Duan, Q.; Wang, Z.; Koplev, S.; Jenkins, S.L.; Jagodnik, K.M.; Lachmann, A.;, et al. Enrichr: a comprehensive gene set enrichment analysis web server 2016 update. Nucleic Acids Res 2016, 44, W90–97, doi:10.1093/nar/gkw377.

64. Lambert, S.A.; Jolma, A.; Campitelli, L.F.; Das, P.K.; Yin, Y.; Albu, M.; Chen, X.; Taipale, J.; Hughes, T.R.; Weirauch, M.T. The Human Transcription Factors. Cell 2018, 172, 650–665, doi:10.1016/j.cell.2018.01.029.

65. Xu, W.; Ye, Y.; Sharrocks, A.D.; Zhang, W.; Chen, X. Genome-wide Interrogation of Protein-DNA Interactions in Mammalian Cells Using ChIPmentation. STAR Protoc 2020, 1, 100187, doi:10.1016/j.xpro.2020.100187.

66. Langmead, B.; Salzberg, S.L. Fast gapped-read alignment with Bowtie 2. Nat Methods 2012, 9, 357–359, doi:10.1038/nmeth.1923.

67. Li, H.; Handsaker, B.; Wysoker, A.; Fennell, T.; Ruan, J.; Homer, N.; Marth, G.; Abecasis, G.; Durbin, R. The Sequence Alignment/Map format and SAMtools. Bioinformatics 2009, 25, 2078–2079, doi:10.1093/bioinformatics/btp352.

68. Zhang, Y.; Liu, T.; Meyer, C.A.; Eeckhoute, J.; Johnson, D.S.; Bernstein, B.E.; Nusbaum, C.; Myers, R.M.; Brown, M.; Li, W.; Liu, X.S. Model-based Analysis of ChIP-Seq (MACS). Genome Biology 2008, 9, R137, doi:10.1186/gb-2008-9-9-r137.

69. Zarrineh, P.; Darieva, Z.; Bobola, N. Motif2Site: a Bioconductor package to detect accurate transcription factor binding sites from ChIP-seq. bioRxiv 2022, 2022.2009.2022.509048, doi:10.1101/2022.09.22.509048.

70. Liao, Y.; Smyth, G.K.; Shi, W. The R package Rsubread is easier, faster, cheaper and better for alignment and quantification of RNA sequencing reads. Nucleic Acids Res 2019, 47, e47, doi:10.1093/nar/gkz114.

71. Robinson, M.D.; McCarthy, D.J.; Smyth, G.K. edgeR: a Bioconductor package for differential expression analysis of digital gene expression data. Bioinformatics 2010, 26, 139–140, doi:10.1093/bioinformatics/btp616.

72. Lawrence, M.; Huber, W.; Pagès, H.; Aboyoun, P.; Carlson, M.; Gentleman, R.; Morgan, M.T.; Carey, V.J. Software for computing and annotating genomic ranges. PLoS Comput Biol 2013, 9, e1003118, doi:10.1371/journal.pcbi.1003118.

73. Larsson, J. Venn diagrams with eulerr. Available online: https://cran.r-project.org/web/packages/eulerr/vignettes/venn-diagrams.html (accessed on 28/03/2024).

74. Ross-Innes, C.S.; Stark, R.; Teschendorff, A.E.; Holmes, K.A.; Ali, H.R.; Dunning, M.J.; Brown, G.D.; Gojis, O.; Ellis, I.O.; Green, A.R.;, et al. Differential oestrogen receptor binding is associated with clinical outcome in breast cancer. Nature 2012, 481, 389–393, doi:10.1038/nature10730.

75. Lun, A.T.; McCarthy, D.J.; Marioni, J.C. A step-by-step workflow for low-level analysis of single-cell RNA-seq data with Bioconductor. F1000Res 2016, 5, 2122, doi:10.12688/f1000research.9501.2.

76. Langfelder, P.; Zhang, B.; Horvath, S. Defining clusters from a hierarchical cluster tree: the Dynamic Tree Cut package for R. Bioinformatics 2008, 24, 719–720, doi:10.1093/bioinformatics/btm563.

